# The search for sexually antagonistic genes: Practical insights from studies of local adaptation and statistical genomics

**DOI:** 10.1101/2020.04.30.071191

**Authors:** Filip Ruzicka, Ludovic Dutoit, Peter Czuppon, Crispin Y. Jordan, Xiang-Yi Li, Colin Olito, Homa Papoli Yazdi, Anna Runemark, Erik I. Svensson, Tim Connallon

**Author notes:** Denotes equal contributions (Filip Ruzicka, Ludovic Dutoit).

## Abstract

Sexually antagonistic (SA) genetic variation—in which genotypes favoured in one sex are disfavoured in the other—is predicted to be common and has been documented in several animal and plant populations, yet we currently know little about its pervasiveness among species or its population genetic basis. Recent applications of genomics in studies of SA genetic variation have highlighted considerable methodological challenges to the identification and characterisation of SA genes, raising questions about the feasibility of genomic approaches for inferring SA selection. The related fields of local adaptation and statistical genomics have previously dealt with similar challenges, and lessons from these disciplines can therefore help overcome current difficulties in applying genomics to study SA genetic variation. Here, we integrate theoretical and analytical concepts from local adaptation and statistical genomics research—including *F*_*ST*_ and *F*_*IS*_ statistics, genome-wide association studies (GWAS), pedigree analyses, reciprocal transplant studies, and evolve-and-resequence (E&R) experiments—to evaluate methods for identifying SA genes and genome-wide signals of SA genetic variation. We begin by developing theoretical models for between-sex *F*_*ST*_ and *F*_*IS*_, including explicit null distributions for each statistic, and using them to critically evaluate putative signals of sex-specific selection in previously published datasets. We then highlight new statistics that address some of the limitations of *F*_*ST*_ and *F*_*IS*_, along with applications of more direct approaches for characterising SA genetic variation, which incorporate explicit fitness measurements. We finish by presenting practical guidelines for the validation and evolutionary analysis of candidate SA genes and discussing promising empirical systems for future work.

**Impact Summary:** Genome sequences carry a record of the evolutionary and demographic histories of natural populations. Research over the last two decades has dramatically improved our ability to detect genomic signals of adaptation by natural selection, including several widely-used methods for identifying genes underlying local adaptation and quantitative trait variation. Yet the application of these methods to identify sexually antagonistic (SA) genes—wherein variants that are adaptive for one sex are maladaptive for the other—remains under-explored, despite the potential importance of SA selection as a mechanism for maintaining genetic variation. Indeed, several lines of evidence suggest that SA genetic variation is common within animal and plant populations, underscoring the need for analytical methods that can reliably identify SA genes and genomic signals of SA genetic variation. Here, we integrate statistics and experimental designs that were originally developed within the fields of local adaptation and statistical genomics and apply them to the context of sex-specific adaptation and SA genetic variation. First, we evaluate and extend statistical methods for identifying signals of SA variation from genome sequence data alone. We then apply these methods to re-analyse previously published datasets on allele frequency differences between sexes—a putative signal of SA selection. Second, we highlight more direct approaches for identifying SA genetic variation, which utilise experimental evolution and statistical associations between individual genetic variants and fitness. Third, we provide guidelines for the biological validation, evolutionary analysis, and interpretation of candidate SA polymorphisms. By building upon the strong methodological foundations of local adaptation and statistical genomics research, we provide a roadmap for rigorous analyses of genetic data in the context of sex-specific adaptation, thereby facilitating insights into the role and pervasiveness of SA variation in adaptive evolution.

## Introduction

A population’s evolutionary capacity for adaptation hinges upon the nature and extent of the genetic variation it harbours (Fisher 1930). In simple environments where selection is uniform over time, across space, and among different classes of individuals within the population, adaptation may proceed by fixing unconditionally beneficial mutations and eliminating deleterious ones. Yet species exist in complex environments, where the rate of adaptation is constrained by mutations that are favoured in some selective contexts but disfavoured in others. Genetic variants exhibiting such trade-offs respond slowly to natural selection (Otto 2004; Connallon and Hall 2018) and thereby limit the capacity for population persistence (Gomulkiewicz and Houle 2009; Chevin 2013), local adaptation (Lenormand 2002), species’ range expansion (Kirkpatrick and Barton 1997; Duputié et al. 2012), and adaptation in general (Brady et al. 2019).

“Sexually antagonistic” (SA) genetic variation—wherein alleles that are beneficial when expressed in one sex are harmful when expressed in the other—represents a particularly common form of genetic trade-off (Rice and Chippindale 2001; Bonduriansky and Chenoweth 2009; Van Doorn 2009). SA genetic variation arises from sex differences in selection (*a*.*k*.*a*., sex-specific selection) on traits that are genetically correlated between the sexes (Connallon and Clark 2014b), and may contribute substantially to fitness variation (Kidwell et al. 1977; Abbott 2011; Connallon and Clark 2014a; Olito et al. 2018) and maladaptation (Lande 1980; Matthews et al. 2019). Estimates of phenotypic selection suggest that sex differences in directional selection are common (Cox and Calsbeek 2009; Lewis et al. 2011; Gosden et al. 2012; Stearns et al. 2012; Morrissey 2016; De Lisle et al. 2018; Singh and Punzalan 2018), implying that many genetic variants affecting quantitative traits have SA effects on fitness. Likewise, estimates of genetic variation for fitness suggest that the genetic basis of female and male fitness components is partially discordant, with some multi-locus genotypes conferring high fitness in one sex and low fitness in the other (Chippindale et al. 2001; reviewed in Connallon and Matthews 2019).

While studies of sex-specific selection indicate that SA alleles contribute to fitness variation in several animal and plant populations (*e*.*g*., Chippindale et al. 2001; Fedorka and Mousseau 2004; Svensson et al. 2009; Delph et al. 2011; Mokkonen et al. 2011; Berger et al. 2014), the population genetic basis of this fitness variation is largely unknown, leaving several important questions unanswered. For example, what fraction of genetic variance for fitness is attributable to SA alleles versus other classes of genetic variation (*e*.*g*., deleterious mutations)? Is SA genetic variation attributable to many small-effect loci or to a few large-effect loci? Are SA polymorphisms maintained under balancing selection, or are they transient and primarily evolving via mutation, directional selection and drift? Are SA alleles randomly distributed across the genome or are they enriched on certain chromosome types (*e*.*g*., sex chromosomes)? These questions are part of the broader debate about the genetic basis of fitness variation and the evolutionary forces that maintain it (Lewontin 1974; Charlesworth and Hughes 1999), which is one of the oldest in this field and perhaps the most difficult to resolve (*e*.*g*., Lewontin 1974, p. 23).

As in most areas of evolutionary biology, research on SA selection is increasingly drawing upon genomics. A few studies have identified candidate SA polymorphisms with large effects on traits related to fitness (Roberts et al. 2009; Barson et al. 2015; Rostant et al. 2015; VanKuren and Long 2018; Pearse et al. 2019), and this handful of SA loci almost certainly represents the tip of the iceberg (Ruzicka et al. 2019). Many other studies have highlighted genomic patterns of expression or sequence diversity that could be indicative of sex-specific selection (*e*.*g*., Innocenti and Morrow 2010; reviewed in Kasimatis et al. 2017; Mank 2017; Rowe et al. 2018).

While there is optimism that genomics will facilitate the study of sex-specific selection, we still face several challenges in applying genomic data to identify and characterise SA genetic variation. For example, some putative genomic signals of sex-specific selection, such as sex-biased gene expression, are ambiguous: at best, they may serve as indirect proxies of sex-specific selection, or at worst provide no information about contemporary selection on each sex (Kasimatis et al. 2017; Rowe et al. 2018). Allele frequency differences between sexes (*e*.*g*., between-sex *F*_*ST*_) may represent more straightforward genomic signatures of sex-specific selection (*e*.*g*., Lucotte et al. 2016; Cheng and Kirkpatrick 2016). Yet ambiguous null hypotheses for empirical estimates of between-sex *F*_*ST*_, along with high statistical noise relative to biological signal in these estimates, raise questions about statistical power and the prevalence of false positives within such data (Kasimatis et al. 2019). In addition, we need to better understand the extent to which common pitfalls of genome sequence datasets—*e*.*g*., mis-mapped reads, sampling biases, hidden population structure, and impacts of linkage and hitchhiking—yield artefactual signals of sex-specific processes (*e*.*g*., Bissegger et al. 2019; Kasimatis et al. 2020), and thus questionable candidate SA genes.

The challenges in applying genomics to sex-specific selection have strong parallels in the fields of local adaptation and statistical genomics. However, whereas the study of SA loci is still in its infancy, the fields of local adaptation and statistical genomics have already grappled intensively with many of the conceptual and methodological challenges that research on sex-specific selection now faces (Hoban et al. 2016; Visscher et al. 2017). For example, local adaptation research has long emphasized the importance of clear null models for distinguishing genes involved in local adaptation from false positives that simply reside in the tails of neutral null distributions (Lewontin and Krakauer 1973; Günther and Coop 2013; Whitlock and Lotterhos 2015). Similarly, statistical genomics researchers have repeatedly warned that hidden population structure in genome-wide association studies (GWAS) can lead to spurious conclusions about the genetic basis of quantitative traits, complex diseases, and the role of adaptation in population differentiation (Lander and Schork 1994; Price et al. 2010; Barton et al. 2019; Berg et al. 2019; Sohail et al. 2019). Lessons from local adaptation and statistical genomics research can therefore sharpen hypothesis framing, guide statistical methodology, and inform best practices for disentangling signal, noise, and artefacts in studies of sex-specific selection.

Here, by drawing insights from local adaptation and statistical genomics research, we present practical guidelines for population genomic analyses of sex-specific fitness variation. We first outline two statistics that can, in principle, provide *indirect* evidence of sex-specific fitness effects of genetic variation: between-sex *F*_*ST*_, which is sensitive to sex differences in viability selection and some components of reproductive success, and *F*_*IS*_, a measure of Hardy-Weinberg deviations in diploids, which is sensitive to sex differences in overall selection (*i*.*e*., cumulative effects of viability, fertility, fecundity and mating competition).

We develop theoretical null models for each metric, provide an overview of their sampling distributions and statistical power, and present a reanalysis of published *F*_*ST*_ data in light of our models. We also highlight complementary methods adapted from case-control GWAS to overcome some of the limitations of these metrics. Second, we evaluate several *direct* approaches for characterising sex-specific genetic variation for fitness, which combine elements from quantitative genetics, reciprocal transplant studies and experimental evolution. These direct approaches have been extensively employed to study the genetic basis of locally adapted phenotypes and quantitative traits, but rarely to identify SA loci. Third, we outline approaches that can support the biological validity of candidate genes and discuss best practices for the analysis and interpretation of their evolutionary histories.

### I. Indirect approaches for identifying SA genes

Estimating fitness under natural conditions is difficult, rendering approaches for identifying SA genes that rely on fitness measurements (*i*.*e*., direct approaches; see Section II) unfeasible for many populations. Any widely applicable approach must therefore make use of *indirect* empirical signals of SA selection in genome sequence data, which can now be collected for virtually any species.

Two specific patterns of genome sequence variation could be indicative of contemporary SA selection, as emphasized by several recent studies (*e*.*g*., Cheng and Kirkpatrick 2016; Lucotte et al. 2016; Eyer et al. 2019). First, because sex differences in selection give rise to allele frequency differences between females and males of a population (Box 1), large allele frequency differences between *samples* of females and males of a population could be indicative of sex differences in selection (including SA selection or ongoing sexually concordant selection that differs in magnitude between sexes). Ideally, inferences of sex-specific selection from allele frequency estimates should be based on samples of breeding adults that have passed the filter of viability selection, sexual selection, and fertility/fecundity selection, though in practice genome sequences of random samples of adults are more readily obtainable and will reflect pre-adult viability selection (Kasimatis et al. 2019). Second, elevated allele frequency differences between breeding females and males of a given generation generate elevated heterozygosity in offspring of the next generation relative to Hardy-Weinberg expectations (Kasimatis et al. 2019). Inflated heterozygosity in a large random sample of offspring could therefore reflect sex differences in viability selection, sexual selection, and/or fertility and fecundity selection during the previous generation.

#### Single-locus signals of sex-specific selection in fixation indices

Fixation indices, which are widely applied in studies of population structure (Wright 1951), can be used to quantify allele frequency differences between sexes (*F*_*ST*_) and elevations in heterozygosity in the offspring of a given generation (*F*_*IS*_), each of which are predicted consequences of SA selection (Boxes 1-2). Several studies have estimated *F*_*ST*_ between sexes using gene sequences sampled from adults (Cheng and Kirkpatrick 2016; Lucotte et al. 2016; Flanagan and Jones 2017; Wright et al. 2018, 2019; Bissegger et al. 2019) or from breeding individuals (Dutoit et al. 2018), yet it remains unclear how much information about SA selection is contained within these estimates. A major problem, as emphasized by Kasimatis et al. (2019; see also: Cheng and Kirkpatrick 2016; Kasimatis et al. 2017; Connallon and Hall 2018), is that the effects of sex-specific selection on between-sex *F*_*ST*_ are expected to be weak relative to effects of random sampling variability. Genuine signal of SA selection in the distribution of *F*_*ST*_ estimates may therefore be swamped by sampling error.

As we show in Appendix A (see Box 2), a null distribution for between-sex *F*_*ST*_ estimates at loci with no sex differences in selection conforms to a special case of Lewontin and Krakauer’s (1973) classic null model for *F*_*ST*_ estimated between populations. An appealing feature of our two-sex null model is its insensitivity to some of the simplifying assumptions inherent in Lewontin and Krakauer’s original model (*i*.*e*., that subpopulations are independent; see Robertson 1975; Nei and Maruyama 1975; Charlesworth 1998; Beaumont 2005; Whitlock and Lotterhos 2015), or issues arising from genetic linkage (Charlesworth 1998), which do not affect the two-sex null distribution when *F*_*ST*_ is independently estimated per SNP, but can strongly impact the null for concatenated sequences (see Booker et al. 2020; Appendix A).

By comparing the distribution of between-sex *F*_*ST*_ under the null with the distribution under SA selection (Box 2), we can formally evaluate the minimum strength of selection (*s*_min_) required for a SA locus to reliably reside within the upper tail of the null distribution. For example, the 99^th^ percentile for the null distribution is 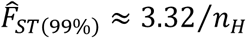, where *n*_*H*_ is the harmonic mean sample size of female- and male-derived gene sequences, and 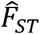 refers to an estimate of *F*_*ST*_ (Appendix A); roughly 1% of *F*_*ST*_ estimates should fall above this threshold when there are no sex differences in selection. The probability that a SA locus resides within the tail of the null distribution depends on the allele frequencies at the locus, the strength of selection, and the sample size of individuals that are sequenced (Appendix A). In studies with large sample sizes (*i*.*e*., *n*_*H*_ = 10^5^ or greater, as in some human genomic datasets: see Fig. 1), 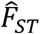 for a SA locus with intermediate allele frequency and fitness effect of a few percent will reliably fall in the upper tail of the null distribution (Fig. 1). In contrast, studies where *n*_*H*_ < 10^4^ require very strong selection for SA loci to reliably reside within the upper tail of the null *F*_*ST*_ distribution (Fig. 1), and are unlikely to identify individual SA loci (*i*.*e*., significant *F*_*ST*_ outliers), even in cases where SA genetic polymorphism *is* common across the genome.

**Figure 1.**
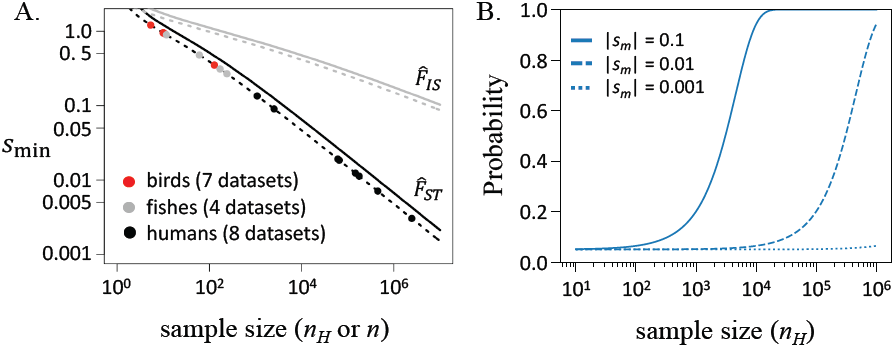
Signal of SA selection relative to sampling error in *F*_*ST*_ and *F*_*IS*_ estimates. **Panel A:** Lines show the minimum strength of selection (*s*_min_; see Appendix A & B) that causes the expected values of 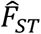 and 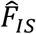 at an SA locus to reach the 5% tail (broken line) or the 1% tail (solid line) of the null distributions for 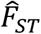 (black lines) and 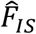 (grey lines). Circles show the theoretical *s*_min_ values (for the 5% tail) for published between-sex *F*_*ST*_ studies that include eight human datasets (Cheng and Kirkpatrick 2016; Lucotte et al. 2016; Kasimatis et al. 2020; Pirastu et al. 2020), four fish datasets (see Flanagan and Jones 2017; Wright et al. 2018, 2019; Bissegger et al. 2019; Vaux et al. 2019), and seven bird datasets (see Dutoit et al. 2018; Wright et al. 2019). **Panel B:** Probability that 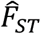 for an additive SA locus (*i*.*e*., *h*_*f*_ = *h*_*m*_ = ½) with intermediate equilibrium allele frequencies (*p, q* = 1/2) is in the top 5% tail of the null distribution for 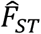. |*s*_*m*_| is the fitness cost of being homozygous for the “wrong” SA allele (see Appendix A & B).

A second putative signal of SA selection is an enrichment of heterozygotes among offspring cohorts, as inferred from high values of *F*_*IS*_ (see Box 2; Appendix B). While only a single study has used *F*_*IS*_ estimates 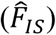 to test for sex-specific selection (Eyer et al. 2019; see Boxes 1-2), the potential for future applications warrants evaluation of signals of SA selection using this metric. Hardy-Weinberg deviations in a sample, as captured by 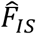, may arise from selection, non-random mating (*e*.*g*., inbreeding or population structure), or random sampling of genotypes from the population (Crow and Kimura 1970; Weir 1997; Lachance 2009). Statistical properties of Hardy-Weinberg deviations in genotype samples are well established (Weir 1997), and easily adaptable for our point of interest: the distribution of 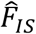 in a randomly mating population in which allele frequencies may differ between the female and male parents of a given generation (Box 2). As illustrated in Box 2, the sampling variance for 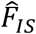 exceeds that of 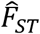 by a substantial margin. Consequently, 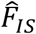 has far less power than 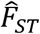 to distinguish signal of sex-specific selection from noise (Fig. 1A).

### *F*_*ST*_ distributions and multi-locus signals of sex-specific selection

Genome scans for individually significant SA loci (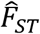 or 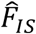 outliers) are severely underpowered unless SA loci segregate for intermediate frequency alleles with large fitness effects and sample sizes are very large (see above). Indeed, no empirical *F*_*ST*_ study to date has yielded individually significant candidate SA SNPs that have survived corrections for multiple-testing and rigorous controls for genotyping error and read-mapping artefacts (see below).

Although *F*_*ST*_ scans for individually significant SA loci are highly conservative, the full empirical distribution of *F*_*ST*_ for autosomal SNPs may nonetheless carry a cumulative signature of SA selection at many loci—even in the absence of individually significant SA genes. For example, SNPs responding to SA selection (*i*.*e*., SA SNPs or those in linkage disequilibrium with them) should inflate the total number of observations in the upper quantiles predicted under the null (Fig. 2). An excess of observations in the upper quantiles of the theoretical null may therefore imply an enrichment of SA SNPs in the tail of the empirical 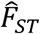 distribution, with SNPs in the tail representing interesting candidates for follow-up analyses (see section III). Moreover, the fraction of true versus false positives among candidates can be quantified. For example, if 2% of observed SNPs fall within the top 1% quantile of the theoretical null, this implies a 1:1 ratio of true to false positives within the top 2% of observations (*i*.*e*., a false discovery rate of 50%).

**Figure 2.**
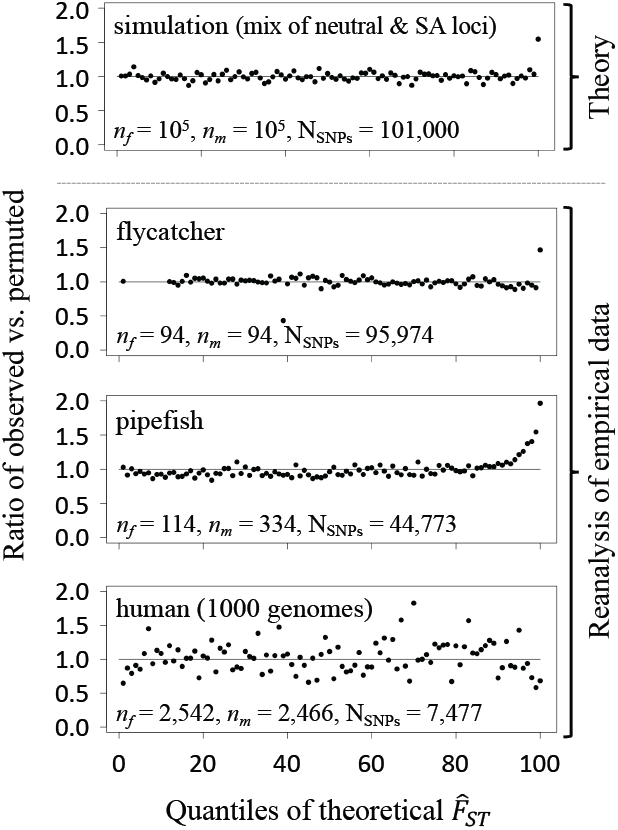
Empirical signals of multi-locus SA polymorphism. Each panel shows the ratio of observed versus permuted 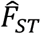 SNPs within 100 quantiles of the theoretical null of 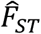. A ratio above one indicates an excess of observed 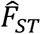 values in a given quantile. **Theoretical data:** 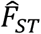 values were simulated for 10^5^ neutrally evolving loci with MAF above 5% in the dataset, and for 103 loci responding to SA selection prior to sampling of gene sequences (each SA-responding locus had a SA fitness effect or was in perfect linkage disequilibrium with an SA locus). SA-responding loci had intermediate allele frequencies in the population (*p* = ½). SA selection coefficients were drawn from an exponential distribution with mean of *s*_*avg*_ = 0.03, with additive fitness effects (*h* = ½). The top 1% theoretical quantile is enriched by ∼50%, implying that ∼1/3 of SNPs above the 99% threshold of the null are true positives. **Empirical data:** population samples from flycatchers, pipefish, and humans (1000 Genomes data), excluding variants with MAF < 0.05; Supplementary Fig. S5 illustrates the effect of including rare variants in the human re-analysis. For flycatcher data, the absence of values in some low-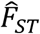 quantiles is due to the small number of sampled sequences, which generates some discrepancy between the *discrete* distribution of 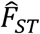 estimates (for both observed and permuted data) and the *continuous* theoretical null outlined in Box 2. The top 1% theoretical quantiles for flycatchers and pipefish are enriched by ∼50% and ∼100%, respectively, implying that ∼1/3 and ∼1/2 of SNPs above the 99% threshold of the null are “true” positives (which still require biological validation and filtering for putative artefacts, as described in the main text).

To test for elevation of empirical 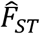 estimates relative to our theoretical null model, we re-analysed three representative datasets from previously published studies (collared flycatcher *Ficedula albicollis*: Dutoit et al. 2018; gulf pipefish *Syngnathus scovelli*: Flanagan and Jones 2017; human: The 1000 Genomes Project Consortium 2015 data used by Cheng and Kirkpatrick 2016). For human and flycatcher whole-genome resequencing datasets, we used autosomal coding variants, excluding any SNP with missing data. For the pipefish RAD-seq dataset, coding and non-coding variants with less than 50% missing data were included; sex-linked regions are unknown in this species and could not be excluded. In all datasets, polymorphic sites with minor allele frequencies (MAFs) less than 5% were also excluded, as sites with low MAF exhibit inflated sampling variances (see Whitlock and Lotterhos 2015; Appendix A and Supplementary Fig. S5). Analyses were carried out in bedtools (Quinlan and Hall 2010), vcftools (Danecek et al. 2011) and R (R Core Team 2020).

For all three datasets, permuted 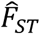 distributions (*i*.*e*., 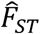 calculated after randomly permuting sex labels across individuals) conform well to the theoretical null model for 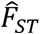 (Supplementary Figs. S1-S4). For the 1000 Genomes human dataset, 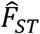 observations (*i*.*e*., non-permuted estimates) are indistinguishable from both the theoretical null and permuted distributions, with no enrichment of observations within the top quantiles of the null (Fig. 2; *χ*^2^) tests; 5% tail: *p* = 0.10; 1% tail: *p* = 0.31). In contrast, flycatcher and pipefish datasets show elevated 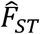 values relative to the 5% and 1% tails of their null distributions (Fig. 2; *χ*^2^) tests; *p* = 4.33 × 10^−9^ and *p* < 2.2 × 10^−16^ for flycatcher, *p* < 2.2 × 10^−16^ and *p* < 2.2 × 10^−16^ for pipefish). Such enrichment could imply that many loci are responding to sex differences in selection (either directly or indirectly through hitchhiking with selected loci), or that a false signal of elevated 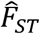 has been generated by population structure and/or data quality issues, as discussed below.

#### Accounting for spurious signals of sex-specific selection

The analyses presented above suggest that 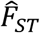, though severely underpowered for detecting individually significant outlier SNPs, may capture signals of SA SNP enrichment within the tail of the 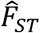 distribution. However, sex differences in selection should not be invoked as the cause of such elevations in 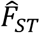 without excluding artefacts that may generate similar patterns.

First, incorrect mapping of sex-linked markers to autosomes can potentially lead to artificial inflation of *F*_*ST*_ estimates. For example, Y- or W-linked sequence may be erroneously mapped to sequence paralogs on autosomes, resulting in artificially high *F*_*ST*_ inferences at autosomal sites (Bissegger et al. 2019; Kasimatis et al. 2020). This problem is difficult to control for in species lacking high-quality reference genomes, including those where the sex determination system is unknown (*e*.*g*., the pipefish dataset in Fig. 2). Moreover, the effects of demographic processes, including recent admixture events or sex-biased migration, play out differently between sex chromosomes and autosomes (Hedrick 2007), so that caution is required in interpreting elevated between-sex *F*_*ST*_ on the X or Z, as has been reported in humans (*e*.*g*., Lucotte et al. 2016).

Second, sex differences in population structure—arising from the broad geographic sampling of individuals or recent migration into a single sampled population—can also generate signals of genetic differentiation between females and males in the absence of sex differences in selection (Box 1). Taxa with broad contemporary distributions (*e*.*g*., humans, *Drosophila*) often show significant genetic differentiation among populations. Uneven or unrepresentative sampling of individuals of each sex from a set of different locations can, by chance, inflate allele frequency differences between the sexes beyond expectations for a single, panmictic population. If loci showing high between-sex *F*_*ST*_ also exhibit high between-population *F*_*ST*_, this could be indicative of population structure contributing to allele frequency divergence between the sexes in the empirical sample. Studies that sample individuals from a single population may also show artificially elevated between-sex *F*_*ST*_ if migration is sex-biased (Box 1), which is common among animals (Trochet et al. 2016).

Sex-specific population structure can be accounted for by leveraging the statistical framework of case-control GWAS, in which associations between polymorphic variants and binary phenotypic states (*e*.*g*., presence or absence of a disease) are quantified. The case-control GWAS approach would treat *sex* (female or male) as the binary phenotypic state and scan for loci with the strongest associations, which should exhibit elevated absolute odds ratios (see Appendix C; Kasimatis et al. 2020; Pirastu et al. 2020). While the underlying logic is identical to between-sex *F*_*ST*_, existing analytical methods for case-control GWAS can take population structure and relatedness in the empirical sample into account by including kinships (or the top principal components derived from kinships) as covariates (Astle and Balding 2009; Price et al. 2010). The case-control GWAS framework also permits estimation of SNP-based heritability (Yang et al. 2011; Speed et al. 2017) of the phenotype (*i*.*e*., sex), which can be used to test for a genome-wide signal of sex-specific selection.

Despite the advantages of leveraging an existing statistical framework, using case-control GWAS to test for associations between alternative alleles and sex does not sidestep all of the challenges faced by *F*_*ST*_ and *F*_*IS*_ statistics (see Appendix C). As with *F*_*ST*_ and *F*_*IS*_, large sample sizes remain essential for discriminating between sampling variance and true signal of sex differences in selection (especially when selection is weak), and the methods perform poorly when MAFs are low. Additionally, association tests using odds ratios depend heavily on a normal approximation for which the underlying assumptions are often violated, and there is a deep and still evolving literature regarding hypothesis testing using these methods that users should be aware of (*e*.*g*., Haldane 1956; Wang and Shan 2015).

### II. Direct approaches for identifying SA genes

In exceptional study systems, candidate SA polymorphisms can be identified through explicit statistical associations between genotypes and fitness. Such *direct* inference approaches present two major advantages over indirect methods. First, the inclusion of fitness measurements can potentially increase power to detect individual SA loci, relative to indirect methods (*e*.*g*., Fig. 1). Second, association tests can be conducted across many components of fitness (*e*.*g*., viability, fecundity, mating success), facilitating identification of the life history stages and selective contexts affected by SA loci. We outline two general approaches for direct inference of sex-specific selection—genome-wide association studies (GWAS) and evolve-and-resequence (E&R) studies—which have been extensively employed to identify genes associated with human trait variation and/or local adaptation (Long et al. 2015; Visscher et al. 2017), yet rarely to identify SA loci.

#### GWAS of sex-specific fitness

GWAS quantify statistical associations between phenotypic variation and polymorphic SNPs throughout the genome. Using GWAS to identify SA loci further requires that data on fitness components and genotypes are collected from individuals of each sex. A major advantage of GWAS is the availability of statistically rigorous methods to identify candidate loci, including methods to control for covariates in analyses (Price et al. 2010), and approaches that correct for multiple testing (*e*.*g*., family-wise or false discovery rate correction; Benjamini and Hochberg 1995) or that reduce the number of tests through gene-based association analysis (Nagamine et al. 2012; Riggio et al. 2013; Bérénos et al. 2015). We discuss the application of GWAS to three types of dataset: (i) datasets in which genotypes and phenotypes are each measured independently in each sex (*e*.*g*., humans); (ii) systems amenable to experimental manipulation, in which each genotype can be replicated among female and male carriers (*e*.*g*., isogenic or hemi-clone fruit fly lines); (iii) pedigreed populations, in which the genealogical relationships between all individuals are known (*e*.*g*., some sedentary vertebrate populations).

Where genotypic and phenotypic measurements are performed among independently sampled individuals of each sex, as in humans, SA loci can be identified by first performing a separate GWAS in each sex (‘sex-stratified’ GWAS) (Fig. 3A), and then quantifying the difference between male- and female-specific effect sizes (see also Gilks et al. 2014). Illustrating this approach, Winkler et al. (2015) performed a sex-stratified GWAS on several human anthropometric traits, then defined a *t*-statistic as 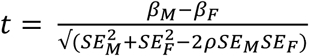, where *β*_*M*_ and *β*_*F*_ are the sex-specific effect sizes, *SE*_*M*_ and *SE*_*F*_ are the sex-specific standard errors, and *ρ* is the between-sex rank correlation among genome-wide loci. For each polymorphic site, *p*-values were generated by comparing the observed *t* statistics to a null *t*-distribution with no sex-specific effects (where E[*t*] = E[*β*_0_ − *β*_1_] = 0 under the null). This approach has been applied to non-fitness traits in humans (Randall et al. 2013; Myers et al. 2014; Winkler et al. 2015; Mitra et al. 2016), but has yet to be applied to fitness components (*e*.*g*., ‘number of children’ phenotype in the UK Biobank; Sudlow et al. 2015).

**Figure 3.**
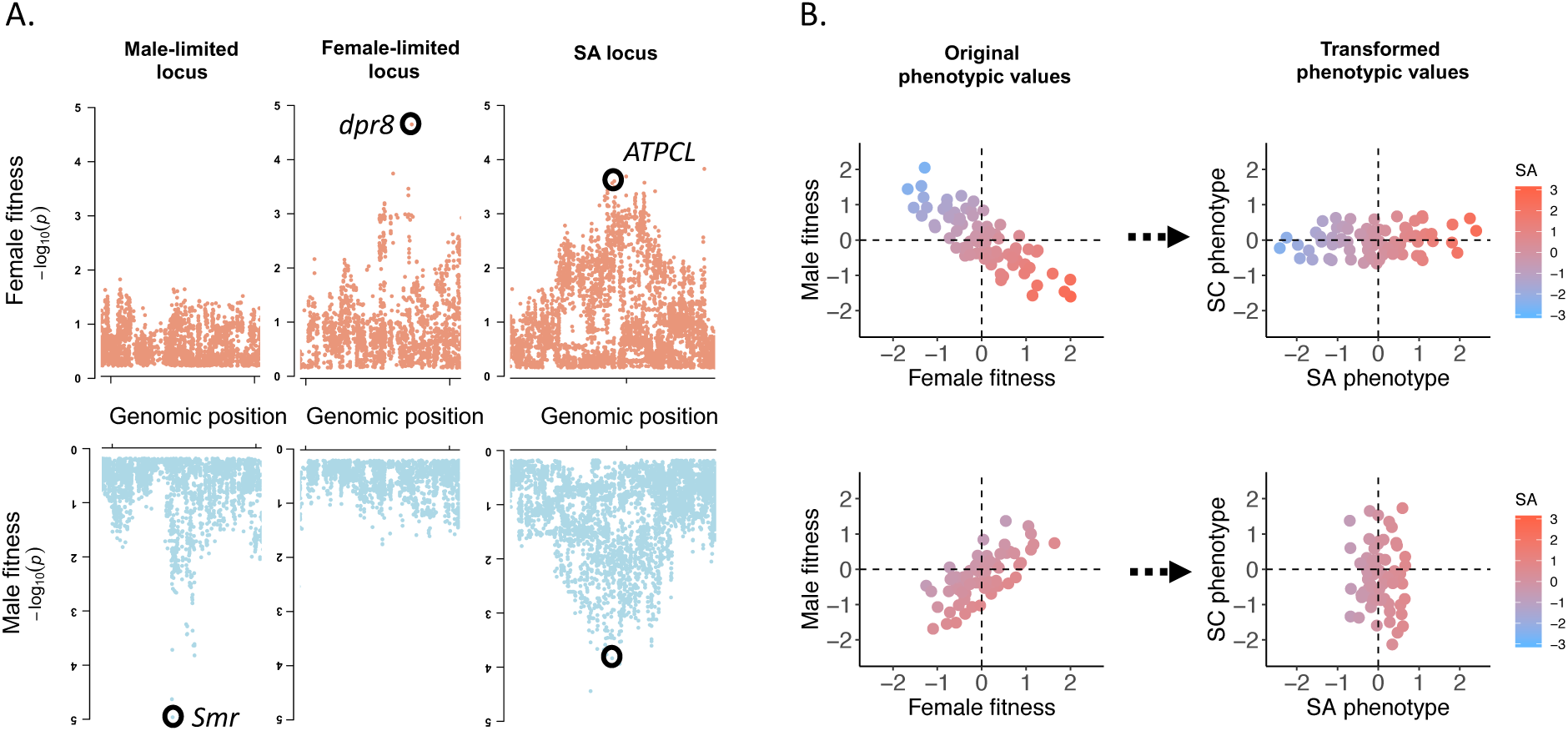
GWAS as a direct method to identify SA loci. **Panel A:** Manhattan plots of –log_10_ P-values from a GWAS of female and male fitness (data from Ruzicka et al. 2019), illustrating distributions of P-values at or near candidate loci (circled) with male-limited (left), female-limited (middle) and SA (right) fitness effects, respectively. **Panel B:** Simulated phenotypic values of female and male fitness, before (left) and after (right) 45-degree rotation of the bivariate coordinate matrix to obtain sexually antagonistic (SA) and sexually concordant (SC) phenotypic values. Where fitness variation is predominantly SA (top left), most variation is along the SA phenotypic axis (top right). Where fitness variation is predominantly SC (bottom left), most variation is along the SC phenotypic axis (bottom right). In all panels, colours denote SA phenotypic values.

In some experimental systems (*e*.*g*., fruit flies; flowering plants), the creation of isogenic or hemi-clone lines (Abbott and Morrow 2011; Mackay et al. 2012; Berger et al. 2014) allows the same genotypes to be replicated and phenotypically assayed in carriers of each sex. Here, genotypes are effectively transplanted into male and female bodies or “environments”, analogous to the reciprocal transplantation of individuals sampled from different environments in local adaptation studies (Price et al. 2018). Identifying SA loci can then be achieved by transforming the bivariate coordinate system of male and female fitness values of a set of genotypes through matrix rotation (see Berger et al. 2014), which generates a univariate SA phenotype amenable to GWAS analysis (Fig. 3B). The approach is exemplified by a recent study in *D. melanogaster* (Ruzicka et al. 2019), which identified ∼230 candidate SA polymorphic sites.

In pedigreed vertebrate populations, such as Soay sheep (*Ovis ares*) or Florida scrub jays (*Aphelocoma coerulescens*), the genetic relationships between all individuals are known, and transmission of individual alleles across successive generations can be estimated (MacCluer et al. 1986). Because an individual’s genetic contribution to future generations is a genuine representation of its Darwinian fitness, alleles transmitted more frequently by one sex relative to the other represent candidate SA variants. Analyses of pedigreed populations of Florida scrub jays have been used to identify alleles with above-average transmission rates to descendants irrespective of sex (*i*.*e*., unconditionally beneficial alleles; Chen et al. 2019), yet this type of GWAS remains to be used to identify SA loci. It should be noted, however, that many pedigreed populations are necessarily small (given the logistics of monitoring them), which may hinder detection of loci affecting fitness variation.

Though GWAS-based identification of candidate SA loci shows promise, two major drawbacks must be kept in mind. First, measurements that capture total lifetime reproductive success are difficult to obtain, and caution is required in interpreting results based on single fitness component (*e*.*g*., reproductive but not viability selection), which may correlate imperfectly with total fitness. Second, GWAS effect sizes are typically small for polygenic traits (Visscher et al. 2017), including fitness. Powerful GWAS of sex-specific fitness may therefore be logistically prohibitive, and candidate SA loci will necessarily represent the subset of loci with particularly large fitness effects.

#### Evolve & Resequence (E&R) with sex-limited selection

Experimental elimination of selection in one sex but not the other (*i*.*e*., sex-limited selection) is a powerful way to identify SA selection in action. Various sex-limited selection designs have been implemented, including: (i) restricting transmission of the genome to the male line, and thereby removing selection through females (Rice 1996, 1998; Prasad et al. 2007; Bedhomme et al. 2008; Abbott et al. 2010) or vice versa (Rice 1992); (ii) eliminating fitness variance in within one sex but not the other (Rundle et al. 2006; Morrow et al. 2008; Maklakov et al. 2009; Hollis et al. 2014; Immonen et al. 2014; Chenoweth et al. 2015; Veltsos et al. 2017); and (iii) applying sex-limited artificial selection on a specific fitness component, such as mating success (Dugand et al. 2019), lifespan (Berg and Maklakov 2012; Chen and Maklakov 2014; Berger et al. 2016), mating type (Bielak et al. 2014), or mating investment (Pick et al. 2017).

To identify SA loci, sex-limited selection can be combined with genotyping at multiple time points during experimental evolution within each selection regime (E&R), thereby connecting population genetic changes to the phenotypic responses accrued during experimental evolution. By tracking allele frequencies in male-limited and female-limited selection lines, alleles that show a significant time-by-treatment interaction point to candidate SA loci, and their frequency dynamics can be characterised using current analytical tools for E&R experiments (Wiberg et al. 2017; Vlachos et al. 2019). E&R is a powerful and proven approach for identifying the genomic basis of phenotypic variation and local adaptation (Turner et al. 2011; Long et al. 2015; Barghi et al. 2019). Moreover, because it is experimental, the issues of sex-specific population structure that arise in between-sex *F*_*ST*_ studies (see section I) can be minimised. Yet despite these advantages, we are not aware of any published study that has used E&R to identify SA loci (see Chenoweth et al. 2015 for the closest effort to date).

E&R can be performed using population samples taken at multiple time-points within a single generation (Svensson et al. 2018) or across multiple generations, with the latter approach benefitting from the fact that allele frequency responses to selection are cumulative over multiple generations. E&R, like GWAS, remains best suited for detecting loci with relatively large fitness effects. Selection on complex polygenic traits typically leads to small changes in allele frequencies at large numbers of loci, resulting in genomic signals of selection that are difficult to distinguish from genetic drift (Schlötterer et al. 2015). Consequently, the study organism, the number of replicates, the effective population sizes of selection and control lines, and the duration of experiments must be carefully considered in the design of E&R experiments (Baldwin-Brown et al. 2014; Kofler and Schlötterer 2014; Kessner and Novembre 2015).

### III. Validation and follow-up analyses of candidate SA genes

Candidate SA genes and SNP sets enriched for SA alleles (*i*.*e*., identified using methods outlined above) provide context for addressing long-standing questions about SA variation, including the genomic distribution, biological functions, and population genetic processes shaping SA polymorphisms. We focus on two specific issues in follow-up analyses of putative SA variants. First, we outline approaches for biologically validating SA candidates—a crucial task given that candidate gene sets may include appreciable proportions of false positives and artefactual signals of sex-specific selection (see section I). Second, we discuss population genetic analyses and issues of interpretation with bearing on the evolutionary histories of SA genes.

#### Biological validation of SA genes

Candidate SA loci can be directly validated in laboratory-amenable taxa by experimentally manipulating each allele and measuring its sex-specific fitness effect. A good example of experimental validation of naturally occurring SA genes is a study by VanKuren and Long (2018), in which RNA interference and CRISPR-Cas9 were used to demonstrate SA effects of tandem duplicate genes *Apollo* and *Artemis* on offspring production in *D. melanogaster*. Similarly, Akagi and Charlesworth (2019) used manipulative molecular experiments to study candidate SA genes in several plant species. As a third example, several studies investigated a P450 transposable element insertion that upregulates the *Cyp6g1* gene and increases DDT resistance in *D. melanogaster* (Smith et al. 2011; Rostant et al. 2015, Hawkes et al. 2016). Though evidence for SA effects at this particular locus is mixed, the experimental approaches used—including measurements of sex-specific fitness among isogenic lines and tracking the frequencies of each alternative allele in experimental cages—represent validation steps with potential for broad usage.

Direct experimental manipulation of candidate SA genes is not always feasible, and in instances where it is not, their biological validity can be assessed in other ways. One way is to test whether candidate loci are enriched in genomic regions that are putatively functional (*e*.*g*., coding or regulatory) rather than inert (*e*.*g*., intergenic). Such “genic enrichment”, which is expected for SA polymorphisms with genuine phenotypic effects, has previously been used to strengthen validity of candidate alleles for local adaptation (Barreiro et al. 2008; Coop et al. 2009; Key et al. 2016). Another way to increase confidence in candidate loci is to look for multiple signals of SA selection. For example, candidates identified through elevated *F*_*ST*_ that also exhibit elevated heterozygosity (high *F*_*IS*_), or are associated with SA fitness effects in a GWAS, represent the best candidates for follow-up evolutionary analyses (see below). Finally, if independent data exist on the sex-specific phenotypic effects of individual mutations (*e*.*g*., in RNAi databases), these data can be mined to support the validity of candidate SA genes.

#### Evolutionary dynamics of SA genes

We do not outline the range of population genetic analyses that could be used to describe the evolutionary dynamics of candidate SA loci, as these have been comprehensively reviewed elsewhere (*e*.*g*., Vitti et al. 2013; Fijarczyk and Babik 2015). Instead we provide guidance on some common issues that are likely to arise when analysing patterns of genetic variation at SA loci and interpreting their mode of evolution.

First, we emphasise that the evolutionary dynamics of a contemporary SA gene may have, in the past, been governed by any combination of genetic drift, net directional selection (selection favouring fixation of one SA allele), or balancing selection (selection maintaining SA polymorphism). Theory often focuses on the conditions generating balancing selection at SA loci (*e*.*g*., Kidwell et al. 1977; Patten and Haig 2009; Fry 2010), leading some empirical studies to use signals of balancing selection (*e*.*g*., elevated Tajima’s D) as indirect proxies for SA selection (*e*.*g*., Dutoit et al. 2018; Wright et al. 2018, 2019; Sayadi et al. 2019). However, whether or not contemporary candidate SA alleles evolved under balancing selection hinges upon both the historical pattern of sex-specific selection and dominance at such loci (Kidwell et al. 1977; Connallon and Chenoweth 2019) and the effective population size (Connallon and Clark 2012, 2014a; Mullon et al. 2012). SA loci with large and symmetric selection coefficients or beneficial reversals of dominance (*e*.*g*., *h*_*f*_ = 1 and *h*_*m*_ = 0 in Box 1) are most conducive to balancing selection, whereas sufficient asymmetry between the sexes in the strength of selection (*e*.*g*., Mallet and Chippindale 2011; Mallet et al. 2011; Sharp and Agrawal 2013) should result in net directional selection that removes SA polymorphism rather than maintaining it (Kidwell et al. 1977). Even when conditions for long-term balancing selection are met, the *efficacy* of balancing selection relative to drift may often be weak at SA loci, leading to genetic diversity patterns that are indistinguishable from neutrally evolving loci (Connallon and Clark 2012, 2013; Mullon et al. 2012). In short, loci under contemporary SA selection can have a broad range of possible evolutionary histories. As such, the typical mode of evolution operating at candidate SA loci *cannot be assumed a priori* and should instead be viewed as a question that must be resolved empirically.

Second, the detection of elevated polymorphism at SA loci does not necessarily imply balancing selection. For example, SA candidate loci may exhibit significantly elevated MAFs relative to non-SA loci (Ruzicka et al. 2019), yet relaxed directional selection can account for this pattern if non-SA loci encompass a mix of neutral sites and sites evolving under sexually concordant directional selection. To establish that SA loci are evolving under balancing selection, it is necessary to show that SA genetic variation is significantly elevated compared to confirmed neutral sites (*e*.*g*., short introns, Parsch et al. 2010) and cannot be accounted for by demographic or mutational processes (Andrés et al. 2009; DeGiorgio et al. 2014; Bitarello et al. 2018). On the other hand, significant *reductions* in polymorphism at SA loci, relative to neutral sites, do not necessarily rule out balancing selection either. Counterintuitively, when the equilibrium frequency of the minor allele is rare (*i*.*e*., equilibrium MAF < 0.2, approximately), balanced polymorphisms can be lost more rapidly than neutral polymorphisms, leading to reduced genetic variation relative to neutral expectations (Robertson 1962; Mullon et al. 2012).

A third and final point is that non-random patterns of genetic variation at SA loci can be generated by ascertainment bias alone. For example, data filtering steps that remove low-MAF SNPs (see section I) necessarily exclude rare SA variants from all downstream analyses. Elevated power to detect fitness effects among intermediate-frequency sites in a GWAS can also generate a spurious positive relationship between candidate SA sites and MAF that might be mistaken for non-neutral evolution. Similarly, SA loci could be non-randomly distributed across the genome (*e*.*g*., among hypermutable CpG sites, or regions of elevated recombination), thereby generating spurious patterns in population genomic data that appear to indicate non-neutral evolution. It is therefore important to correct for such biases where possible by, for example, incorporating external data on recombination rate variation (Comeron et al. 2012; Elyashiv et al. 2016) or assessing evidence of trans-species polymorphisms among SA loci before and after removal of CpG sites (Leffler et al. 2013).

### IV. Moving Forward

We have critically assessed a broad range of methods for detecting genomic signatures of SA selection, including *indirect methods* based on genome sequence analysis (section I), and *direct methods* based on associations between genome sequences and fitness measurements (section II). An inescapable conclusion from our indirect inference models (*i*.*e*., *F*_*ST*_, *F*_*IS*_, and odds-ratio statistics) is that very strong sex differences in selection or very large sample sizes are required to detect individual SA candidate polymorphisms with high confidence (Fig. 1), in agreement with previous simulation studies of between-sex *F*_*ST*_ (Lucotte et al. 2016; Connallon and Hall 2018; Kasimatis et al. 2019). However, estimates of the full distribution of *F*_*ST*_ from previously published flycatcher and pipefish datasets reveal an intriguing elevation of genome-wide *F*_*ST*_, relative to our null models, that justifies future empirical studies of allele frequency differences between sexes (see below). While direct methods for identifying SA genes must circumvent the substantial logistical challenge of accurately measuring fitness, the approach is powerful when feasible (see Ruzicka et al. 2019) and certain to be a key component of future work on the genetics of sex-specific fitness variation.

While there is little doubt that identifying and characterising SA genes is challenging, there are several reasons for optimism. First, the low power of indirect metrics to detect selection at an individual-locus level does not rule out the detection of a cumulative signal of polygenic sex differences in selection (*e*.*g*., Fig. 2). Although such an approach implies that candidate SA genes (*e*.*g*., those in the highest *F*_*ST*_ quantiles) will include many false positives, elevated false discovery rates are not necessarily problematic if we are interested in the general properties of SA candidates relative to samples of putatively neutral (or non-SA) loci. Nevertheless, in studies with low-to-moderate sample sizes, where many candidate genes will be false positives, researchers should minimally demonstrate that: (i) the empirical distribution of the metric of interest differs significantly from its appropriate null (see Kasimatis et al. 2019; our re-analyses in section I), (ii) putative signals of selection are not driven by sex-specific population structure or other artefacts (see section I), and (iii) candidate loci are biologically plausible (see section III).

Second, the power to detect SA genes using indirect metrics can often be increased in relatively simple ways. For example, pooled sequencing is a cost-effective method for estimating allele frequencies from samples of many individuals (Schlötterer et al. 2015), and well-suited for genome-wide *F*_*ST*_ scans. Researchers could, alternatively, focus attention towards large publicly available genomic datasets that are adequately powered for detecting loci under moderately strong SA selection (see Fig. 1), or towards genomic regions predicted to have relatively high statistical power. For example, studies of pseudo-autosomal regions of recombining sex chromosomes have substantially higher power to detect *F*_*ST*_ outliers driven by sex differences in selection (Qiu et al. 2013; Kirkpatrick and Guerrero 2014). Creative sampling strategies may also amplify power to identify SA genes. For example, estimating allele frequencies among *breeding* adults—which have passed filters of viability selection and components of adult reproductive success—increases the number of episodes of selection that can contribute to allele frequency differentiation between sexes, improving the potential for detecting elevated between-sex *F*_*ST*_.

Third, well-chosen study systems can improve prospects for accurately measuring lifetime reproductive success and identifying SA loci through direct methods (GWAS or E&R). For example, difficulties in accurately measuring fitness under field conditions can be mitigated in pedigreed populations, where the genetic contribution of each individual to successive generations is known (provided the population is well monitored), and each genotype can therefore be associated with an accurate estimate of total lifetime reproductive success in each sex. Emerging approaches to infer pedigrees from genomic data alone (Snyder-Mackler et al. 2016) may further facilitate identification of SA loci in the absence of long-term monitoring efforts. In some experimental systems, such as laboratory-adapted hemiclones of *D. melanogaster* (Rice et al. 2005; Abbott and Morrow 2011), relatively accurate measurements of outbred lifetime reproductive success are also possible. E&R is feasible for experimental organisms with short generation times and where large laboratory populations can be maintained (*e*.*g*., *Drosophila*, seed beetle *Callosobruchus maculatus*).

Here there is a relatively untapped opportunity to identify SA loci by combining sex-limited selection (*e*.*g*., Rice 1992; Prasad et al. 2007; Morrow et al. 2008; Abbott et al. 2010; Bonel et al. 2018) with genotyping across multiple generations (*e*.*g*., Turner et al. 2011; Long et al. 2015; Barghi et al. 2019; Abbott et al. 2020).

Finally, despite notable exceptions (*e*.*g*., the dioecious plant *Silene latifolia*; Muyle et al. 2012; Delph et al. 2011), plant systems remain underutilised in research on SA selection. One advantage of plants is their greater amenability to field measurements of fitness components, as widely utilized in studies of local adaptation and species’ range limits (Hargreaves et al. 2014). Another advantage is the great diversity of reproductive systems in flowering plant species, the vast majority of which are hermaphroditic and susceptible to SA selection (Jordan and Connallon 2014; Tazzyman and Abbott 2015; Olito 2017; Olito et al. 2018), potentially leading to allele frequency differences between the pollen and ovules contributing to fertilization, and to elevated *F*_*IS*_ among offspring. A third advantage of plants is their greater tendency to express genetic variation during the haploid stage of their life cycle (*e*.*g*., Immler and Otto 2018). Haploid (relative to diploid) expression is expected to inflate the contribution of genetic polymorphism to fitness variance and magnify evolutionary responses to selection, including within-generation allele frequency divergence between sexes (Connallon and Jordan 2016). Exploiting plant systems may thereby increase statistical power to identify candidate SA genes or genomic signals of SA variation using direct (GWAS, E&R) or indirect inference approaches.

## Acknowledgments

We thank Alison Wright, Hans Ellegren and Sarah Flanagan for sharing data, Sarah Flanagan for support through pipefish data analysis, and Isobel Booksmythe for helpful comments on an earlier draft of this manuscript. The study was supported by funds from the Australian Research Council to T.C. and F.R., The Swedish Research Council (VR: grant number 2016-03356) to E.I.S., and the Swiss National Science Foundation (Ambizione number 180145) to X.Y.L. The New Zealand eScience Infrastructure (NeSI) provided support through their high-performance computing facilities; NeSI and collaborator institutions are funded through the Ministry of Business, Innovation & Employment’s Research Infrastructure programme (https://www.nesi.org.nz). We are particularly grateful to the European Society for Evolutionary Biology (ESEB) for funding the Special Topics Network workshops ‘Linking local adaptation with the evolution of sex differences’, which motivated and enabled this study.

## Box 1.

**Processes generating sexually divergent allele frequencies**

Several evolutionary scenarios can lead to sex differences in the frequencies with which individual alleles are transmitted to offspring (Hedrick 2007; Úbeda et al. 2011; Connallon et al. 2018). We focus on two processes—sex differences in selection and sex-biased migration—that may each commonly arise and affect estimates of allele frequency differences between sexes.

### Sex differences in selection

Consider a single bi-allelic locus in which the focal allele (allele *A*) has a frequency of *p* at birth within a given generation; the alternative *a* allele has a frequency of *q* = 1 – *p*. Selection during the life cycle alters the allele frequencies in the set of breeding adults contributing to offspring of the next generation. The frequency of the *A* allele in breeding females and males (respectively) is:

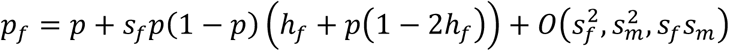

and

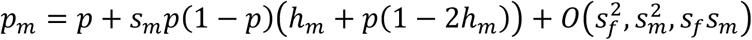

where *s*_*f*_ and *s*_*m*_ are female and male selection coefficients for the *A* allele, *h*_*f*_ and *h*_*m*_ are the dominance coefficients (Table 1), and 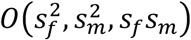 refers to second-order terms in the selection coefficients, which are negligible (and can be ignored) when *s*_*f*_ and *s*_*m*_ are small, as expected for most loci (Charlesworth and Charlesworth (2010), p. 97).

**Table 1.** Sex-specific relative fitness of genotypes of a bi-allelic locus

|  | **Genotype** |  |  |
| --- | --- | --- | --- |
|  | *AA* | *Aa* | *aa* |
| female relative fitness | $1 + s_f$ | $1 + s_f h_f$ | 1 |
| male relative fitness | $1 + s_m$ | $1 + s_m h_m$ | 1 |

We expect allele frequency differences between breeding adults of each sex (*p*_*f*_ ≠ *p*_*m*_) when the fitness effects of each allele differ between sexes, *i*.*e*.: (1) the same allele is favoured in each sex but the strength of selection differs between sexes (*e*.*g*., *p*_*f*_ > *p*_*m*_ when *s*_*f*_ > *s*_*m*_ > 0); or (2) alleles are sexually antagonistic (*e*.*g*., *p*_*f*_ > *p*_*m*_ when *s*_*f*_ > 0 > *s*_*m*_). Allele frequencies are expected to remain equal between sexes (*p*_*f*_ = *p*_*m*_) when genetic variation is neutral (*s*_*f*_ = *s*_*m*_ = 0), or selection and dominance coefficients are identical between sexes (*s*_*f*_ = *s*_*m*_ and *h*_*f*_ = *h*_*m*_).

### Sex-biased migration

Consider an island population receiving new migrants each generation, with migration occurring before reproduction during the life cycle. At birth, the frequencies of *A* and *a* alleles in the island population are *p* and *q* = 1 – *p*, respectively. Let *m*_*f*_ and *m*_*m*_ represent the proportions of breeding females and breeding males on the island that are migrants. The expected frequency of the *A* allele in breeding females and males (respectively) will be:

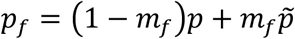

and

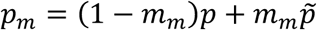

where 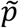 is the frequency of the *A* allele in migrant individuals. The identity, 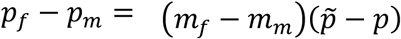, implies that allele frequency differences between breeding females and males (*p*_*f*_ ≠ *p*_*m*_) would require sex-biased migration (*m*_*f*_ ≠ *m*_*m*_) *and* allele frequency differences between migrant and resident (non-migrant) individuals 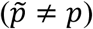.

## Box 2.

**Fixation indices (*F***_***ST***_ **and *F***_***IS***_**) applied to sex differences**

Allele frequency differences between breeding adults of each sex can be quantified by way of fixation indices, originally devised by Wright (1951) for characterising genetic differentiation among populations. We again consider a bi-allelic autosomal locus with the focal allele (*A*) at a frequency of *p*_*f*_ in breeding females and *p*_*m*_ in breeding males.

### Between-sex F_ST_

*F*_*ST*_ is a standardized measure of the allele frequency difference between the sexes:

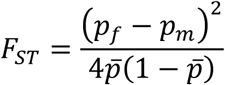

where 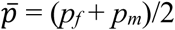 (Cheng and Kirkpatrick 2016). This definition applies to the entire population, and therefore differs from empirical estimates of *F*_*ST*_ that are based upon samples of gene sequences from the population. Though there are several estimators of *F*_*ST*_ (Bhatia et al. 2013), we focus on the simplest:

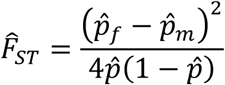

where 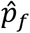 and 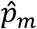 are the allele frequency estimates from samples of females and males, and 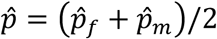. As we show in the Appendix, the ratio

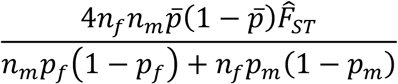

has an approximately noncentral chi-squared distribution with one degree of freedom and noncentrality parameter of 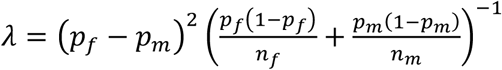, where *n*_*f*_ and *n*_*m*_ are the numbers of sequences derived from females and males, respectively. The approximation can break down when *n*_*f*_ and *n*_*m*_ are small or the minor (rarer) allele at the locus has a frequency near to zero. Under the statistical null distribution, in which the true allele frequencies do not differ between the sexes, 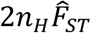 has a chi-squared distribution with one degree of freedom, where *n*_*H*_ = 2(1/*n*_*f*_ + 1/*n*_*m*_)^−1^ is the harmonic mean sample size.

### F_IS_ in offspring

*F*_*IS*_ can be used to quantify deviations between the observed heterozygosity in offspring and the expected heterozygosity under Hardy-Weinberg equilibrium. With random mating among the breeding adults, and ignoring effects of genetic drift or segregation distortion, the frequency of the *A* allele in offspring will be 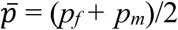, and the proportion of offspring that are heterozygous will be *P*_*Aa*_ = *p*_*f*_(1 – *p*_*m*_) + *p*_*m*_(1 – *p*_*f*_). Under these conditions, *F*_*IS*_ will be:

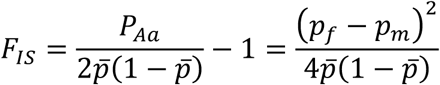

where the final expression is equivalent to *F*_*ST*_ between breeding females and males of the prior generation (Kasimatis et al. 2019). The above expression for *F*_*IS*_ applies to the entire set of offspring in a population, whereas empirical estimates of *F*_*IS*_ will be based on the genotypes of offspring sampled from the population. As shown in the Appendix, estimates of *F*_*IS*_ will be approximately normally distributed with a mean and variance of:

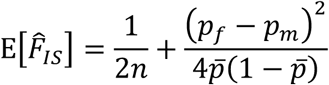

and

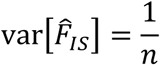

where *n* is the number of offspring sequenced for the locus. The approximation applies when the sample size is large. Under a null model, in which offspring are outbred and mating is random, estimates 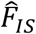 will be normal with mean of 1/2*n* and variance of 1/*n*.

