## Supplementary Material for "The search for sexually antagonistic genes: Practical insights from studies of local adaptation and statistical genomics"

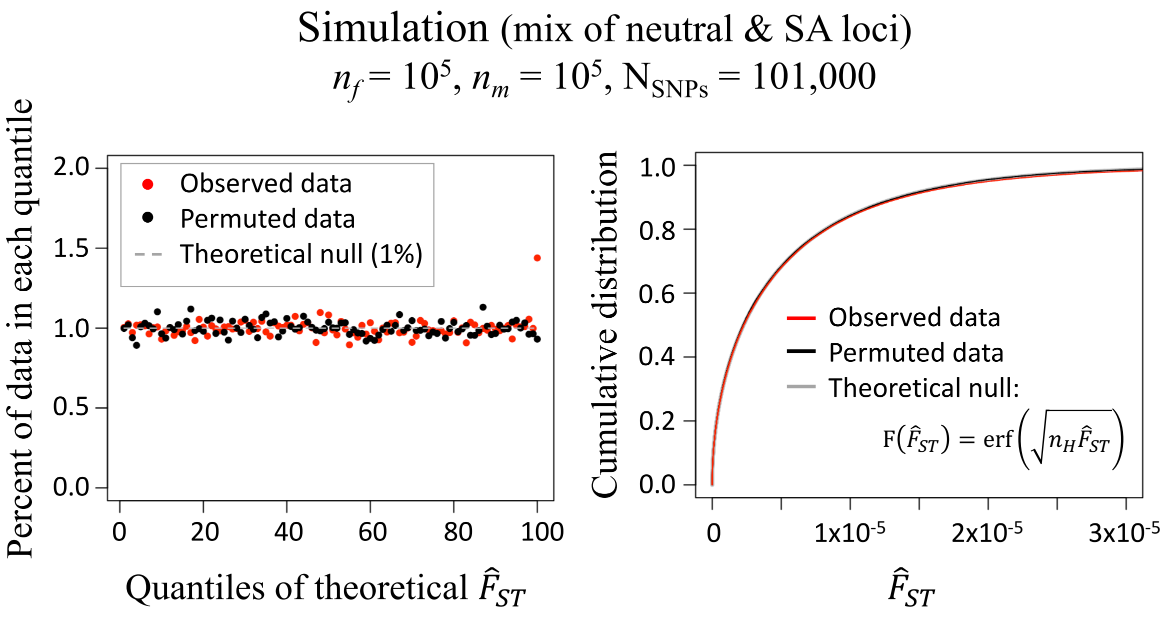


**Figure S1.** Permuted vs. observed $\hat{F}_{ST}$ for simulated data. Details follow those described in Fig. 2 of the main text.

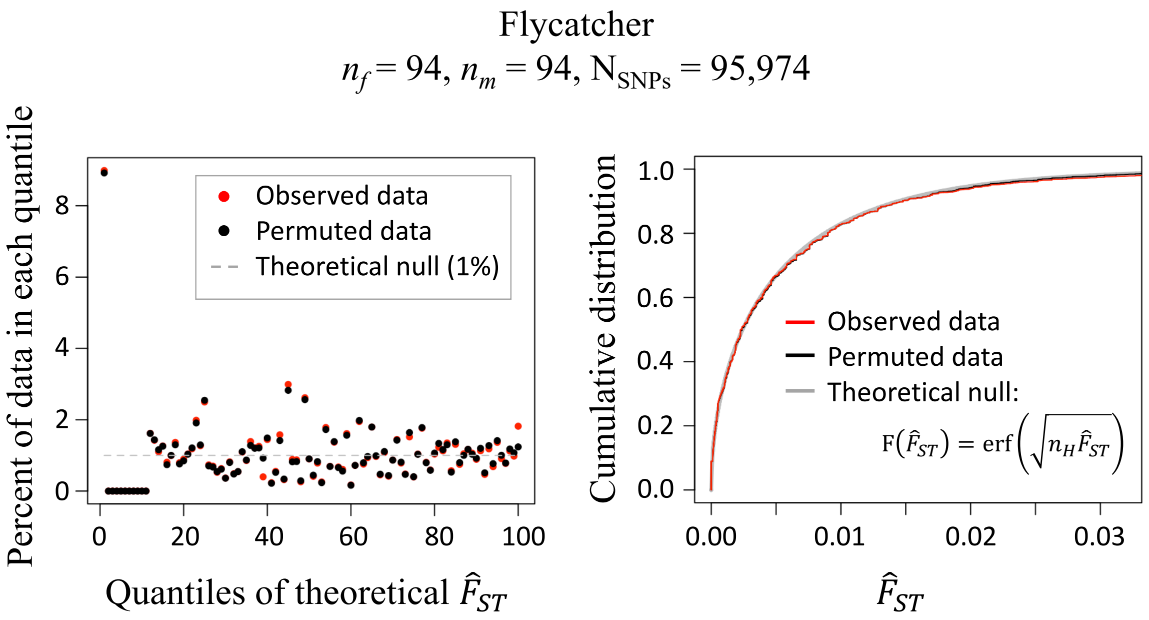


**Figure S2.** Permuted vs. observed $\hat{F}_{ST}$ for the flycatcher dataset. Details of the analysis follow those described in Fig. 2 of the main text.

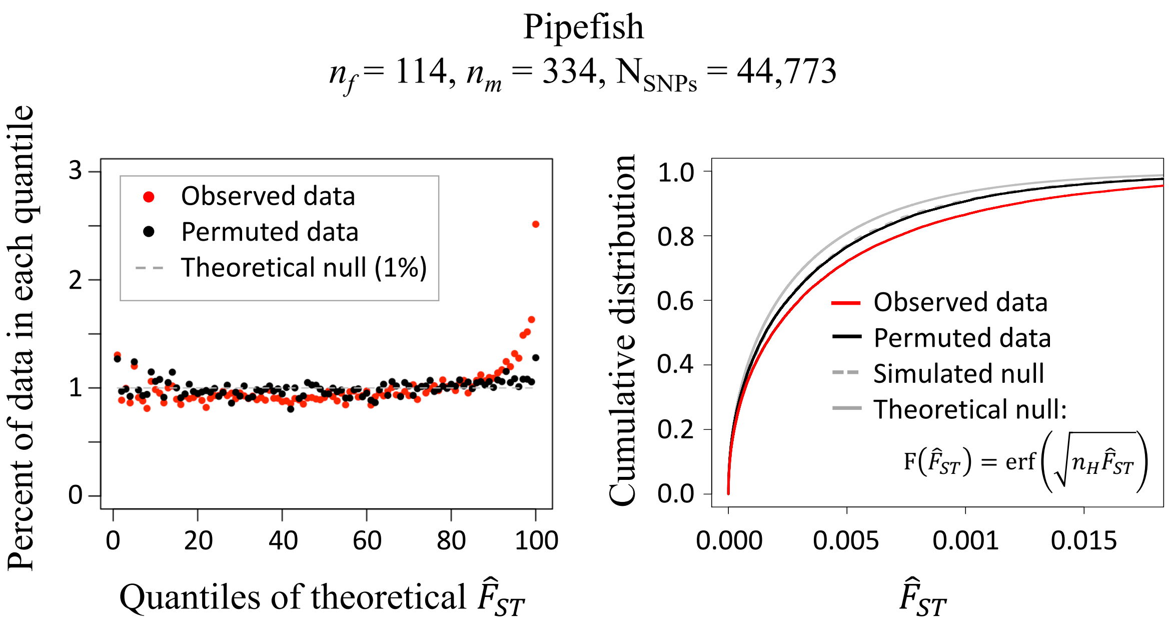


**Figure S3.** Permuted vs. observed $\hat{F}_{ST}$ for the pipefish dataset. Details of the analysis follow those described in Fig. 2 of the main text. Note that there are two null distributions in the right-hand panel. One is the idealized theoretical null distribution (the solid grey curve) which would apply if there was no variation in sample sizes across SNPs (*i.e.*, it requires that $n_H$ is roughly constant across polymorphic sites in the genome). The idealized null model is inappropriate for the pipefish dataset due to missing data leading to sample size variation among sites. The simulated null accounts for missing data by simulating $\hat{F}_{ST}$ per site from a null distribution with site-specific $n_H$.

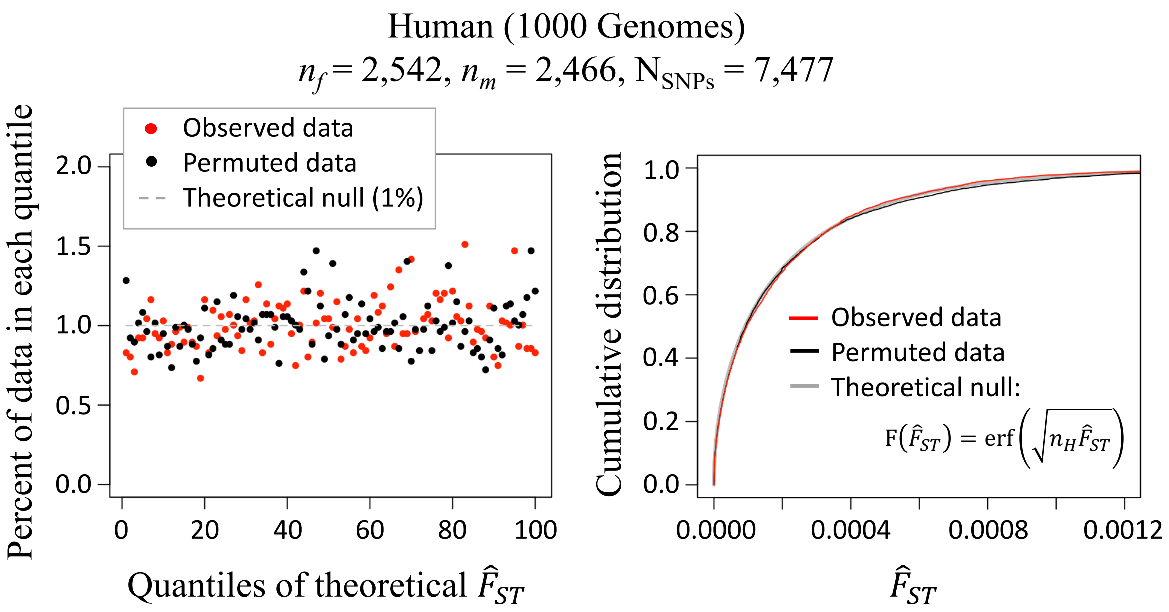


**Figure S4.** Permuted vs. observed $\hat{F}_{ST}$ for the human 1000 Genomes dataset. Details of the analysis follow those described in Fig. 2 of the main text.

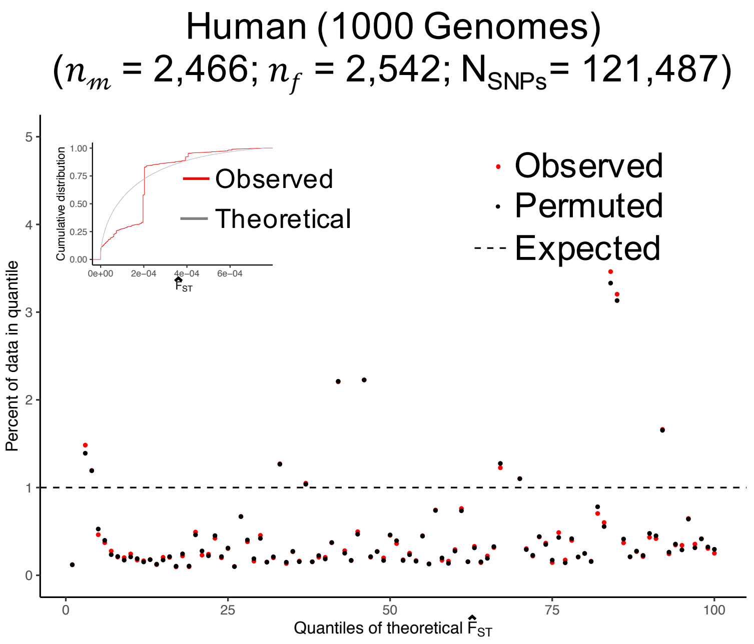


**Figure S5:** Human 1000 Genomes data with rare MAFs included in the analysis. All else is the same as Fig. S4.

### Appendix A: Distribution of $F_{ST}$ estimates

Following Cheng and Kirkpatrick (2016), between-sex $F_{ST}$ for a bi-allelic locus is defined as:

$$F_{ST} = \frac{\left(p_f - p_m\right)^2}{4\bar{p}(1-\bar{p})}$$

where $p_f$ and $p_m$ are the female and male population frequencies of the focal allele at the locus, and $\bar{p}$ is the sex-averaged allele frequency, $\bar{p} = \left(p_f + p_m\right)/2$. The above expression applies to the entire population. Define $F_{ST}$ estimated from allele frequencies in a sample of gene sequences as:

$$\hat{F}_{ST} = \frac{\left(\hat{p}_f - \hat{p}_m\right)^2}{4\hat{p}(1-\hat{p})}$$

where $\hat{p}_f$ and $\hat{p}_m$ are the female and male frequencies calculated from the sample of sequences, and $\hat{p} = \left(\hat{p}_f + \hat{p}_m\right)/2$. We note that several other $F_{ST}$ estimators are widely used in the empirical literature (Bhatia et al 2013), and we return to these further below.

Suppose that, at birth, the focal allele at a given locus has an overall population frequency of $p$ in both sexes (the other allele at the locus has a frequency of $1-p$). Although genetic drift can lead to different allele frequencies between the sexes at birth, we assume that the population size is sufficiently large that such differences are negligible. As long as the sample size of sequenced individuals is large and the strength of selection at the locus is not extreme, then the allele frequencies in the panel of sequenced adults will be similar to the allele frequencies at birth in the population. We can therefore approximate $F_{ST}$ from the sample of sequences using a Taylor Series expansion around $\hat{p}_f = \hat{p}_m = p$:

$$\begin{aligned}\hat{F}_{ST} = {} & \left(\hat{p}_f - p\right)\frac{\partial \hat{F}_{ST}(\hat{p}_i = p)}{\partial \hat{p}_f} + (\hat{p}_m - p)\frac{\partial \hat{F}_{ST}(\hat{p}_i = p)}{\partial \hat{p}_m} \\ & + \frac{1}{2}\left[\left(\hat{p}_f - p\right)^2 \frac{\partial^2 \hat{F}_{ST}(\hat{p}_i = p)}{\partial {\hat{p}_f}^2} + (\hat{p}_m - p)^2 \frac{\partial^2 \hat{F}_{ST}(\hat{p}_i = p)}{\partial {\hat{p}_m}^2} \right. \\ & \left. + 2\left(\hat{p}_f - p\right)(\hat{p}_m - p)\frac{\partial^2 \hat{F}_{ST}(\hat{p}_i = p)}{\partial \hat{p}_f \partial \hat{p}_m}\right] + O[(\hat{p}_i - p)^3] \approx \frac{\left(\hat{p}_f - \hat{p}_m\right)^2}{4p(1-p)}\end{aligned}$$

with the final approximation dropping third-order and higher terms of $\hat{p}_i - p$.

Given the overall allele frequencies of $p_f$ and $p_m$ in adult females and males of the population, then $n_f \hat{p}_f$ and $n_m \hat{p}_m$ from samples of gene sequences will be independent, binomial distributed random variables: $\hat{p}_i \sim B(n_i, p_i)/n_i$, where $n_i$ is the number of sequences derived from members of the $i$th sex. With sufficiently large $n_i$, the distribution of $\hat{p}_i$ will be approximately normal, $\hat{p}_i \sim N\left(n_i p_i, n_i p_i (1 - p_i)\right)/n_i$, leading to:

$$\hat{p}_f - \hat{p}_m \sim N\left(\left(p_f - p_m\right), \left(\frac{p_f\left(1-p_f\right)}{n_f} + \frac{p_m(1-p_m)}{n_m}\right)\right)$$

Using the normal approximation, $\hat{F}_{ST}$ simplifies to:

$$\hat{F}_{ST} \approx \frac{\left(\frac{p_f(1-p_f)}{n_f} + \frac{p_m(1-p_m)}{n_m}\right)}{4p(1-p)} \left(\frac{\hat{p}_f - \hat{p}_m}{\sqrt{\frac{p_f(1-p_f)}{n_f} + \frac{p_m(1-p_m)}{n_m}}}\right)^2$$

$$= \frac{n_m p_f(1-p_f) + n_f p_m(1-p_m)}{4n_f n_m p(1-p)} X$$

where $X$ is a non-central chi-squared distributed random variable with one degree of freedom, and a mean and variance of $\mathrm{E}[X] = 1 + (p_f - p_m)^2 \left(\frac{p_f(1-p_f)}{n_f} + \frac{p_m(1-p_m)}{n_m}\right)^{-1}$ and $\mathrm{var}[X] = 2\left(1 + 2(p_f - p_m)^2 \left(\frac{p_f(1-p_f)}{n_f} + \frac{p_m(1-p_m)}{n_m}\right)^{-1}\right)$. The mean and variance for $\hat{F}_{ST}$ (respectively) are:

$$\mathrm{E}[\hat{F}_{ST}] \approx \frac{1}{4p(1-p)}\left(\frac{p_f(1-p_f)}{n_f} + \frac{p_m(1-p_m)}{n_m} + (p_f - p_m)^2\right)$$

and

$$\mathrm{var}[\hat{F}_{ST}] \approx 2\left(\frac{1}{4p(1-p)}\right)^2 \left(\frac{p_f(1-p_f)}{n_f} + \frac{p_m(1-p_m)}{n_m}\right)\left(\frac{p_f(1-p_f)}{n_f} + \frac{p_m(1-p_m)}{n_m} + 2(p_f - p_m)^2\right)$$

***For a neutral locus*** ($p \approx p_f \approx p_m$), $\hat{F}_{ST}$ reduces to:

$$\hat{F}_{ST} \approx \frac{n_m + n_f}{4n_f n_m} X_0 = \frac{1}{2n_H} X_0$$

where $X_0$ is a chi-squared random variable with one degree of freedom; $n_H = 2/(1/n_f + 1/n_m)$ is the harmonic mean of the sex-specific sample sizes. The mean and variance for $\hat{F}_{ST}$ become:

$$\mathrm{E}[\hat{F}_{ST}] \approx \frac{n_m + n_f}{4n_f n_m} = \frac{1}{2n_H}$$

and

$$\mathrm{var}[\hat{F}_{ST}] \approx \frac{1}{8}\left(\frac{n_m + n_f}{n_f n_m}\right)^2 = \frac{1}{2n_H^2}$$

Under the null model, the variance $\hat{F}_{ST}$ divided by the square of the mean is:

$$\frac{\mathrm{var}[\hat{F}_{ST}]}{\mathrm{E}[\hat{F}_{ST}]^2} \approx 2$$

which is identical to the ratio for a recently subdivided pair of populations (Lewontin and Krakauer 1973; Nei and Chakravarti 1977).

For sufficiently large $n_H$, the median $\hat{F}_{ST}$ for neutral loci is approximately:

$$\mathrm{med}[\hat{F}_{ST}] \approx \frac{1}{2n_H}\left(\frac{7}{9}\right)^3 \approx \frac{0.47}{2n_H}$$

Since the distribution of $\hat{F}_{ST}$ for neutral loci is a simple linear function of chi-squared random variables, we can define the cumulative distribution of neutral $\hat{F}_{ST}$ using known properties of the chi-squared distribution. The cumulative distribution function for the chi-squared distribution with one degree of freedom is:

$$F(\hat{F}_{ST}) = \mathrm{erf}\left(\sqrt{n_H \hat{F}_{ST}}\right)$$

where $\mathrm{erf}(x)$ refers to the error function.

**Effects of selection on allele frequency divergence between sexes**

As in Box 1, supposing that selection is strong relative to genetic drift, then the allele frequency in the *i*th sex, following selection, will be:

$$p_i = p + s_i p(1-p)\big(h_i + p(1-2h_i)\big) + O(s_i^2)$$

where $s_i$ and $h_i$ are the selection and dominance coefficients for the focal allele in the *i*th sex. Under weak selection ($|s_i| << 1$), the first-order approximation (*i.e.*, neglecting $O(s_i^2)$) provides a good approximation for the allele frequencies, per sex, following selection.

The first-order approximation can, however, prove inaccurate in scenarios involving strong selection (*i.e.*, $|s_f|$, $|s_m| > 0.1$), including cases of SA selection with which we are concerned. For the case of arbitrarily strong SA selection, assume that the fitness effects of SA alleles are additive in each sex (*i.e.*, $h_f = h_m = ½$), and the population is at polymorphic equilibrium ($p_f + p_m = 2p$). For this scenario, we can calculate exact allele frequencies per sex, following selection.

Assume that the *A* allele is female-beneficial and the *a* allele is male-beneficial (*i.e.*, $s_f > 0 > s_m$ in Box 1 of the main text). Following Kidwell et al. (1977), we first re-parameterize the general fitness model by substituting $s_m = -|s_m|$, and $t_f = s_f/(1 + s_f)$ so that relative fitnesses in each sex are scaled against the best genotype in that sex (Table S1). With no dominance ($h_f = h_m = ½$), we obtain a symmetric SA parameterization, where $|s_m|$ and $t_f$ represent the fitness costs to females and males (respectively) of carrying the "wrong" allele (Table S1). Both selection parameters are constrained to be between zero and one ($0 < |s_m|$, $t_f < 1$), and $t_f = |s_m|$ represents the special case where the strength of selection is equal between the sexes.

**Table S1.** Sex-specific relative fitness of genotypes of a bi-allelic locus with additive fitness effects in each sex (no dominance; $h_i = ½$)

| | Genotype | | |
|---|---|---|---|
| | ***AA*** | ***Aa*** | ***aa*** |
| **Original parameterization (Box 1)** | | | |
| female relative fitness | $1 + s_f$ | $1 + s_f/2$ | 1 |
| male relative fitness | $1 + s_m$ | $1 + s_m/2$ | 1 |
| **Re-parameterization for a SA locus ($s_f > 0 > s_m$)** | | | |
| female relative fitness | 1 | $1 - t_f/2$ | $1 - t_f$ |
| male relative fitness | $1 - \|s_m\|$ | $1 - \|s_m\|/2$ | 1 |

Given the reparameterization in Table S1, previous theory (see Connallon and Jordan 2016) has shown that the equilibrium *A* allele frequency of a polymorphic SA locus will be:

$$p = \frac{t_f - |s_m| + t_f|s_m|}{2t_f|s_m|}$$

and the equilibrium frequency difference between breeding females and males is:

$$p_f - p_m = \frac{2(1 - p|s_m|)}{|s_m|}\left(\sqrt{1 + \frac{t_f|s_m|p(1-p)}{(1 - p|s_m|)\big(1 - (1-p)t_f\big)}} - 1\right)$$

With an intermediate equilibrium frequency ($p = ½$, which requires that $t_f = |s_m|$), the expected $\hat{F}_{ST}$ for the locus is:

$$\mathrm{E}\big[\hat{F}_{ST}\big] = \frac{1}{2n_H} + \frac{(p_f - p_m)^2}{4p(1-p)}\left(1 - \frac{1}{2n_H}\right)$$

Given the sampling distribution of $\hat{F}_{ST}$ (Box 2), when will SA selection at a locus reliably cause each statistic to reside within the tail of its null distribution? Consider the following approximations for when such outcomes are likely. Assuming that each SA locus has additive fitness effects (*i.e.*, $h_f = h_m = ½$ and $s_f > 0 > s_m$ in Box 1), and each has evolved to an intermediate equilibrium frequency ($p = ½$ and $|s_m| = s_f/(1 + s_f)$, where $|s_m|$ represents the fitness cost to males of inheriting *AA* and $s_f/(1 + s_f)$ is the cost to females of inheriting *aa*; see Connallon and Jordan 2016), $\hat{F}_{ST}$ for a selected locus will have a relatively high probability of residing within the tail of the null when its expected value, $\mathrm{E}\big[\hat{F}_{ST}|SA\big]$, is greater than the percentile that is used to define the tail of the null. Following Box 2, the $k^{\text{th}}$ percentile of the null distribution for $\hat{F}_{ST}$ is:

$$\hat{F}_{ST(k)}|\text{null} = \frac{\left[\text{erf}^{-1}\left(\frac{k}{100}\right)\right]^2}{n_H}$$

where $\mathrm{erf}^{-1}(x)$ refers to the inverse error function. Given the stated assumptions, the expected $\hat{F}_{ST}$ for an SA locus will be greater than the $k^{\mathrm{th}}$ percentile of the null (*i.e.*, $\mathrm{E}\left[\hat{F}_{ST}|SA\right] > \hat{F}_{ST(k)}|\mathrm{null}$), when $|s_m|$ exceeds the following threshold:

$$s_{\min} = \frac{4\sqrt{\dfrac{2\left[\mathrm{erf}^{-1}\left(\frac{k}{100}\right)\right]^2 - 1}{2n_H\left(1-\frac{1}{2n_H}\right)}}}{1 - \dfrac{2\left[\mathrm{erf}^{-1}\left(\frac{k}{100}\right)\right]^2 - 1}{2n_H\left(1-\frac{1}{2n_H}\right)} + 2\sqrt{\dfrac{2\left[\mathrm{erf}^{-1}\left(\frac{k}{100}\right)\right]^2 - 1}{2n_H\left(1-\frac{1}{2n_H}\right)}}}$$

(see Fig. 1A), where $s_{\min}$ represents the minimum strength of selection (*i.e.*, the minimum cost of inheriting the "wrong" SA allele, per sex) satisfying $\mathrm{E}\left[\hat{F}_{ST}|SA\right] \geq \hat{F}_{ST(k)}|\mathrm{null}$ (see the Appendix). Simulations over a range of sample sizes show that $\hat{F}_{ST}$ for a selected locus has a ~40-50% probability of exceeding the 99$^{\mathrm{th}}$ percentile of the null. Even stronger selection is required to satisfy the condition, $\mathrm{E}\left[\hat{F}_{ST}|SA\right] > \hat{F}_{ST(k)}|\mathrm{null}$, when one SA allele at a locus is rarer than the other ($p \neq ½$) and/or dominance is reversed between sexes (*i.e.*, $h_f > ½ > h_m$; see Connallon and Jordan 2016).

**Other $F_{ST}$ estimators**

The $F_{ST}$ estimator presented above and in Box 2 is a biased estimator with an average that is greater than zero under the null model in which there are no allele frequency differences between the sexes in the population as whole (*i.e.*, $\hat{F}_{ST}$, as defined above has an expectation greater than zero when $p_f = p_m$ in the population). Since alternative estimators are often used in empirical studies of population or sex differentiation, it is worth exploring the distribution of $\hat{F}_{ST}$ based on alternative estimators.

The most widely used estimator, Weir-Cockerham's $F_{ST}$ (see Bhatia et al 2013) is defined as:

$$\hat{F}_{ST} = 1 - \frac{\dfrac{n_H}{n_f + n_m - 2}\left[n_f\hat{p}_f\left(1-\hat{p}_f\right) + n_m\hat{p}_m(1-\hat{p}_m)\right]}{\dfrac{n_H}{2}\left(\hat{p}_f - \hat{p}_m\right)^2 + \dfrac{n_H - 1}{n_f + n_m - 2}\left[n_f\hat{p}_f\left(1-\hat{p}_f\right) + n_m\hat{p}_m(1-\hat{p}_m)\right]}$$

where $n_H = 2\left(\frac{1}{n_f} + \frac{1}{n_m}\right)^{-1}$. Assuming large populations and (at most) moderate selection, then the estimator can be approximated using a Taylor Series expansion around $\hat{p}_f = \hat{p}_m = p$, where $p$ is the allele frequency at birth:

$$\begin{aligned}\hat{F}_{ST} &= \frac{1}{1-n_H} + \frac{1}{2}\Bigg[\big(\hat{p}_f - p\big)^2 \frac{\partial^2 F_{ST}(\hat{p}_i = p)}{\partial \hat{p}_f{}^2} + (\hat{p}_m - p)^2 \frac{\partial^2 F_{ST}(\hat{p}_i = p)}{\partial \hat{p}_m{}^2} \\ &\quad + 2\big(\hat{p}_f - p\big)(\hat{p}_m - p)\frac{\partial^2 F_{ST}(\hat{p}_i = p)}{\partial \hat{p}_f \partial \hat{p}_m}\Bigg] + O((\hat{p}_i - p)^3) \\ &\approx \frac{1}{1-n_H} \\ &\quad + \left(\frac{n_H}{n_H - 1}\right)^2 \left(1 - \frac{2}{n_f + n_m}\right) \frac{\big(\hat{p}_f - p\big)^2 + (\hat{p}_m - p)^2 - 2\big(\hat{p}_f - p\big)(\hat{p}_m - p)}{2p(1-p)} \\ &= \frac{1}{1-n_H} + \left(\frac{n_H}{n_H - 1}\right)^2 \left(1 - \frac{2}{n_f + n_m}\right) \frac{\big(\hat{p}_f - \hat{p}_m\big)^2}{2p(1-p)}\end{aligned}$$

Under the neutral null model, we have (approximately):

$$\hat{p}_f - \hat{p}_m \sim N\left(0, p(1-p)\left(\frac{1}{n_f} + \frac{1}{n_m}\right)\right)$$

$$\hat{F}_{ST} \approx \frac{1}{1-n_H} + \frac{n_H}{(1-n_H)^2}\left(1 - \frac{2}{n_f + n_m}\right) X_0$$

where $X_0$ is a chi-squared random variable with one degree of freedom (as above). For large $n_H$, we can approximate the $F_{ST}$ estimator as:

$$\hat{F}_{ST} \approx \frac{1}{n_H}(X_0 - 1)$$

which gives a mean and variance of:

$$\mathrm{E}\big[\hat{F}_{ST}\big] \approx 0$$

$$\mathrm{var}\big[\hat{F}_{ST}\big] \approx \frac{2}{n_H^2}$$

The mean is now centred at zero, which shows that the Weir and Cockerham estimator is approximately unbiased. Nevertheless, the sampling variance remains inversely proportional to the square of the sample size and will therefore show a similar magnitude of sampling variance.

**Genic $F_{ST}$ estimates**

Booker et al. (2020) have recently pointed out that the distribution of $F_{ST}$ estimated from concatenated sequences will strongly depend on the degree of linkage between polymorphic sites in the sequence. Specifically, the chi-square distribution of $\hat{F}_{ST}$ under the null remains applicable when there is no recombination between sites in the concatenated sequence. With recombination, the expected $\hat{F}_{ST}$ remains the same, whereas the variance declines. This effect of concatenation applies to the context of $F_{ST}$ between sexes, as we illustrate below. As such, we advocate using non-concatenated estimates to evaluate fits between observed and theoretical distributions for $\hat{F}_{ST}$.

Using our original $F_{ST}$ estimator, between-sex $F_{ST}$ for a single neutral locus is:

$$\hat{F}_{ST} = \frac{\left(\hat{p}_f - \hat{p}_m\right)^2}{4\hat{p}(1-\hat{p})} = \frac{X_0}{2n_H}$$

where, as before, $X_0$ is a chi-square random variable with one degree of freedom and $n_H$ is the harmonic mean sample size of female and male gene sequences.

Now define a concatenated genic $F_{ST}$ as the average across $k$ polymorphic sites in a given gene. Assuming the $k$ sites are neutral, we have:

$$\hat{F}_{ST} = \frac{1}{k}\sum_{i=1}^{k} \hat{F}_{ST,i} = \frac{1}{k}\sum_{i=1}^{k} \frac{X_i}{2n_{H,i}}$$

where subscripts refer to the $i$th polymorphic site in the gene.

If the $k$ sites are independently segregating and sequence sample sizes are identical across sites ($n_{H,i} = n_H$), then genic $\hat{F}_{ST}$ simplifies to:

$$\hat{F}_{ST} = \frac{1}{2kn_H}\sum_{i=1}^{k} X_i \sim \frac{\chi_k^2}{2kn_H}$$

where $\sum_{i=1}^{k} X_i \sim \chi_k^2$ is a chi-square random variable with $k$ degrees of freedom. The mean and variance for the estimated genic $\hat{F}_{ST}$ are:

$$\mathrm{E}\left[\hat{F}_{ST}\right] = \frac{1}{2n_H}$$

and

$$\mathrm{var}\left[\hat{F}_{ST}\right] = \frac{1}{2kn_H^2}$$

We see that under the null, the expected $\hat{F}_{ST}$ for a SNP and for a genic region will be the same, whereas the former has a $k$-fold greater variance than the latter. Outliers are, therefore, more likely to arise in genes with lower numbers of polymorphisms or nonindependence (high LD) between polymorphic sites.

#### Appendix B: Sex-specific allele frequencies and $F_{IS}$ estimates

We assume an outbred population with random mating between the breeding adults of each sex. Let $p_f$ and $p_m$ represent the allele frequencies in breeding females and males, as in Appendix A. Following Kasimatis et al. (2019), we define $F_{IS}$ as the degree of enrichment in heterozygosity of among offspring of the population, which is a consequence of the allele frequency differences between breeding adults of the prior generation. Defining p as the frequency of the focal allele in offspring of the breeding adults (so that $2p = p_f + p_m$), we have:

$$F_{IS} = \frac{p_f(1-p_m) + p_m(1-p_f)}{2p(1-p)} - 1 = \frac{p_f(1-p_m) + p_m(1-p_f) - 2p(1-p)}{2p(1-p)}$$

$$= \frac{2\left(p - p_f p_m - p(1-p)\right)}{2p(1-p)} = \frac{(p_f - p_m)^2}{4p(1-p)}$$

with the final expression identical to that of $F_{ST}$ for the population (see above). The deviation from Hardy-Weinberg Equilibrium (HWE) can be quantified using a disequilibrium coefficient, $D_A$ (see Weir 1996, p. 94), in which case $F_{IS}$ becomes:

$$F_{IS} = \frac{P_{Aa}}{2p(1-p)} - 1 = \frac{2p(1-p) - 2D_A}{2p(1-p)} - 1 = -\frac{D_A}{p(1-p)}$$

where $P_{Aa}$ is the frequency of heterozygotes in the population, and $D_A = (p(1-p) - P_{Aa}/2)$ quantifies the HWE deviation. Note that $D_A > 0$ implies a deficit of heterozygotes relative to HWE (as expected under inbreeding or population subdivision), whereas $D_A < 0$ implies an excess of heterozygotes (as expected when there are allele frequency differences between the sexes).

##### The null distribution of $F_{IS}$

With large genotype samples (large $n$), empirical estimates of $D_A$, denoted $\widehat{D}_A$, will be approximately normal (Weir 1996, p. 95), with mean and variance:

$$\mathrm{E}\left[\widehat{D}_A\right] = D_A - \frac{p(1-p) + D_A}{2n}$$

$$\mathrm{var}\left[\widehat{D}_A\right] = \frac{p^2(1-p)^2 + (1-2p)^2 D_A - D_A^2}{n}$$

Consequently, $\left(\widehat{D}_A - \mathrm{E}\left[\widehat{D}_A\right]\right)/\sqrt{\mathrm{var}\left[\widehat{D}_A\right]}$ will follow a standard normal distribution with unit variance. If the population really is at HWE (as under the null model), and the sample size is sufficiently large so that the variance in the test statistic can be approximated using $\mathrm{var}\left[\widehat{D}_A\right] = \frac{\hat{p}^2(1-\hat{p})^2}{n}$, then:

$$\frac{\widehat{D}_A}{\sqrt{\mathrm{var}\left[\widehat{D}_A\right]}} = \frac{\widehat{D}_A\sqrt{n}}{\hat{p}(1-\hat{p})} \sim N\left(-\frac{1}{2\sqrt{n}}, 1\right)$$

From this, we see that the estimate of $F_{IS}$ (denoted $\hat{F}_{IS}$) is approximately normal with mean and variance $\mathrm{E}[\hat{F}_{IS}] = 1/2n$ and $\mathrm{var}[\hat{F}_{IS}] = 1/n$, *i.e.*:

$$\hat{F}_{IS} = -\frac{\hat{D}_A}{\hat{p}(1-\hat{p})} = -\frac{1}{\sqrt{n}}\left(\frac{\hat{D}_A\sqrt{n}}{\hat{p}(1-\hat{p})}\right) \sim N\left(\frac{1}{2n}, \frac{1}{n}\right)$$

Simulations of the mean and variance for $\hat{F}_{IS}$ under the null model for $10^5$ polymorphic loci with minor allele frequencies (MAF) greater than 0.05, are plotted below (lines show the analytical predictions, which match up well against the simulated data).

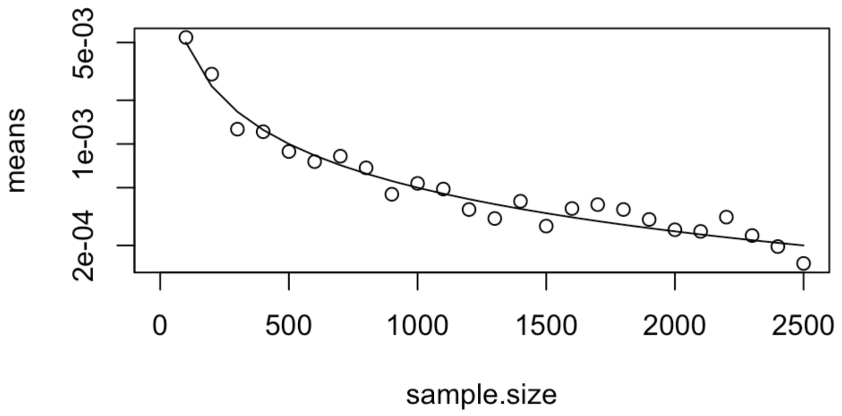


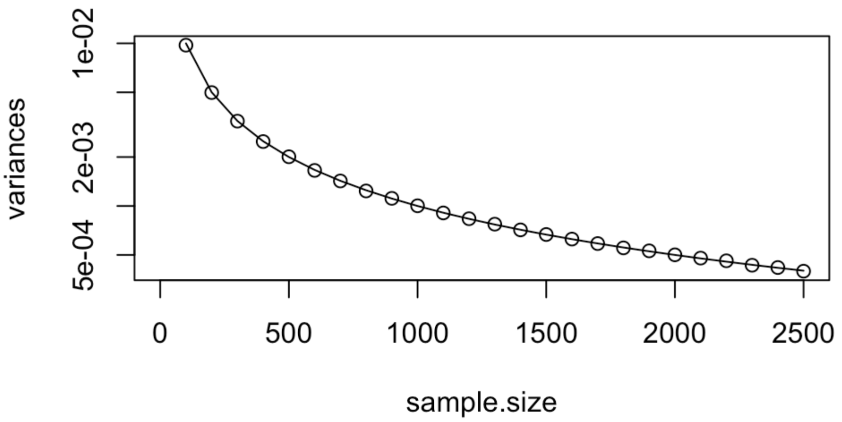


***$F_{IS}$* under sex-specific selection**

With selection, the frequency of heterozygotes at birth will be:

$$p_f(1-p_m) + p_m(1-p_f) = 2p(1-p) + \frac{1}{2}(p_f - p_m)^2$$

where $p = (p_f + p_m)/2$. The HWE deviation in the population becomes:

$$D_A = -\frac{1}{4}(p_f - p_m)^2$$

Assuming that the sample size is large and the HWE deviation is small (|$D_A$| << 1), so that terms of $O(D_A^2, D_A n^{-1})$ can be neglected, the distribution of $\hat{F}_{IS}$ converges, approximately, to:

$$\hat{F}_{IS} = -\frac{\hat{D}_A}{\hat{p}(1-\hat{p})} \sim N\left(\frac{1}{2n} - \frac{D_A}{p(1-p)}, \frac{1}{n}\right)$$

Substituting $D_A = -\frac{1}{4}\left(p_f - p_m\right)^2$, we obtain:

$$\hat{F}_{IS} = -\frac{\hat{D}_A}{\hat{p}(1-\hat{p})} \sim N\left(\frac{1}{2n} + \frac{\left(p_f - p_m\right)^2}{4p(1-p)}, \frac{1}{n}\right)$$

The $k^{\text{th}}$ percentile of the null distribution ($p_f = p_m$) is:

$$\hat{F}_{IS(k)} = \frac{1}{2n} + \sqrt{\frac{2}{n}}\,\mathrm{erf}^{-1}\left(\frac{2k}{100} - 1\right)$$

Using the exact expressions for a polymorphic locus at equilibrium under additive SA selection ($h_i$ = ½), the minimum fitness cost that will cause $\mathrm{E}\left[\hat{F}_{IS}\right]$ to reach the $k^{\text{th}}$ quantile of the null ($\mathrm{E}\left[\hat{F}_{IS}\right] = \hat{F}_{IS(k)}$) is:

$$s_{\min} = \frac{4\sqrt{\sqrt{\frac{2}{n}}\,\mathrm{erf}^{-1}\left(\frac{2k}{100} - 1\right)}}{1 - \sqrt{\frac{2}{n}}\,\mathrm{erf}^{-1}\left(\frac{2k}{100} - 1\right) + 2\sqrt{\sqrt{\frac{2}{n}}\,\mathrm{erf}^{-1}\left(\frac{2k}{100} - 1\right)}}$$

(see Fig. 1A).

### Appendix C: Case-control GWAS and the Log-Odds Ratio

As described in the main text and highlighted in a recent study of SA selection in humans (Kasimatis et al. 2020 bioR$\chi$iv), case-control GWAS methods offer a promising alternative to fixation indices ($F_{ST}$ and $F_{IS}$) for estimating and testing for statistical significance of between-sex allele frequency differences. The advantage of case-control GWAS methods relative to genome-wide scans of between-sex $F_{ST}$ and $F_{IS}$ is that they leverage a well-developed linear modelling framework to explicitly account for confounding factors such as population connectivity when testing for allele frequency differences between sexes. Broadly speaking, case-control GWAS rely on a logistic regression of allele frequency on *sex* (coded as a categorical variable) and other possible explanatory variables, and hypothesis testing is based on the log-odds ratio. Below, we briefly outline the use of the log-odds ratio to test for between-sex differences in allele frequency, illustrate the limits of statistical power when using this approach, and highlight comparisons with the $\hat{F}_{ST}$ statistic, which is presented in the main text.

We emphasize that our brief summary is not meant to be a comprehensive statistical review of odds-ratios, nor a review of case-control GWAS methods and their use to control for population structure and other confounding variables (see *e.g.*, Pirastu et al. 2020; Astle and Balding 2009; Price et al. 2010). We also note that there is a deep and still-evolving statistical literature on hypothesis testing in case-control methods that readers should be aware of (*e.g.*, Haldane 1956, Wang 2015).

**The log-odds ratio ($\mathcal{L}$)**
Given genome sequence data for a random sample of adult males and females, one can construct a 2 × 2 contingency table of allele counts in males and females for a given biallelic locus as:

| | *A* | *a* | *Total* |
|---|---|---|---|
| Males | $n_{m,A}$ | $n_{m,a}$ | $n_m$ |
| Females | $n_{f,A}$ | $n_{f,a}$ | $n_f$ |
| *Total* | $n_A$ | $n_a$ | |

where $n_{i,j}$ is the number of copies of the $j$th allele type in the sample of genes sequenced from the $i$th sex; as in our $F_{ST}$ model $n_f$ and $n_m$ are the total sample sizes of genes sequenced in females and males, respectively. The allele frequency estimates from the samples are:

| | *A* | *a* |
|---|---|---|
| Males | $\hat{p}_{m,A}$ | $\hat{p}_{m,a}$ |
| Females | $\hat{p}_{f,A}$ | $\hat{p}_{f,a}$ |

where $\hat{p}_{i,j} = n_{i,j}/n_i$ with $n_i = n_{i,A} + n_{i,a}$. The sample log-odds ratio can be expressed in the following forms:

$$\hat{\mathcal{L}} = \log\left(\frac{\hat{p}_{m,A}\hat{p}_{f,a}}{\hat{p}_{m,a}\hat{p}_{f,A}}\right) = \log\left(\frac{\hat{p}_{m,A}/\ \hat{p}_{f,A}}{\hat{p}_{m,a}/\ \hat{p}_{f,a}}\right) = \log\left(\frac{\hat{p}_{m,A}}{\hat{p}_{f,A}}\right) - \log\left(\frac{\hat{p}_{m,a}}{\hat{p}_{f,a}}\right)$$

$$= \text{logit}(\hat{p}_{m,A}) - \text{logit}(\hat{p}_{m,a})$$

For large-sample sizes, $\hat{\mathcal{L}}$ is approximately normal with $\hat{\mathcal{L}} \sim \mathrm{N}(\log(\mathrm{OR}), \sigma^2)$, and standard deviation of the gaussian approximation: $SE = \sqrt{\frac{1}{n_{m,A}} + \frac{1}{n_{m,a}} + \frac{1}{n_{f,A}} + \frac{1}{n_{f,a}}}$. We note that hypothesis testing relies heavily on this normal approximation, which breaks down for small sample sizes, and for loci with very low minor allele frequency (MAF) (see *e.g.*, Wang & Shen 2015).

**The null distribution of $\hat{\mathcal{L}}$**

With large genotype samples (large $n = n_f + n_m$), and assuming that there is no sex difference in selection (and thus no allele frequency difference between sexes in the population), empirical estimates of $\hat{\mathcal{L}}$ will be approximately normal with

$$\mathrm{E}[\hat{\mathcal{L}}] = \log\left(\frac{\hat{p}_{m,A}\hat{p}_{f,a}}{\hat{p}_{m,a}\hat{p}_{f,A}}\right)$$

$$var[\hat{\mathcal{L}}] = \frac{1}{n_{m,A}} + \frac{1}{n_{m,a}} + \frac{1}{n_{f,A}} + \frac{1}{n_{f,a}},$$

in which case the scaled log-odds ratio simplifies to $\frac{\hat{\mathcal{L}}}{\sqrt{\frac{1}{n_{m,A}}+\frac{1}{n_{m,a}}+\frac{1}{n_{f,A}}+\frac{1}{n_{f,a}}}} \sim \mathrm{N}(0,1)$, which provides a useful null distribution in the absence of sex-specific selection. The approximation breaks down when any of the allele counts are small, and reliable hypothesis testing therefore requires that the MAF is not too low, or that the sample size is large enough to ensure reasonably high counts of each allele type.

**$\mathcal{L}$ under sex-specific selection**

From Box 1 in the main text, the population frequency of the *A* allele within the *i*th sex, following sex-specific selection, will be:

$$p_i = p + s_i p(1-p)\big(h_i + p(1-2h_i)\big) + O(s_i^2)$$

where $s_i$ and $h_i$ are the selection and dominance coefficients for the focal allele in the *i*th sex. Analytic expressions for $\mathcal{L}$ under sex-specific selection quickly become unwieldy. We can, however, briefly illustrate the expected behavior of the distribution under sex-specific selection in the idealized scenario of intermediate initial allele frequency (*i.e.*, $p$ = ½) and weak and additive sex-specific selection ($h_f = h_m$ = ½; second-order terms $O(s_i^2)$ ignored). Under these simplifying assumptions, the distribution of $\hat{\mathcal{L}}$ is normal with mean and standard deviation:

$$\mathrm{E}[\hat{\mathcal{L}}] = \log\left[\frac{(4-s_f)(4+s_m)}{(4+s_f)(4-s_m)}\right]$$

$$SE = 8\sqrt{\frac{1}{n_f\left(16-s_f^2\right)} + \frac{1}{n_m\left(16-s_m^2\right)}}.$$

Simulations of the mean, standard error, and scaled log odds ratio for $10^4$ loci conforming to the above assumptions show that the approximation works well for an illustrative scenario of SA selection ($s_m$ = 0.03, $s_f$ = -0.03) with equal sample sizes for both sexes (where $i$ indexes sex and $n_m = n_f$), provided sample sizes are sufficiently large. Note that results for the Null in the second panel are overlain by those for SA selection.

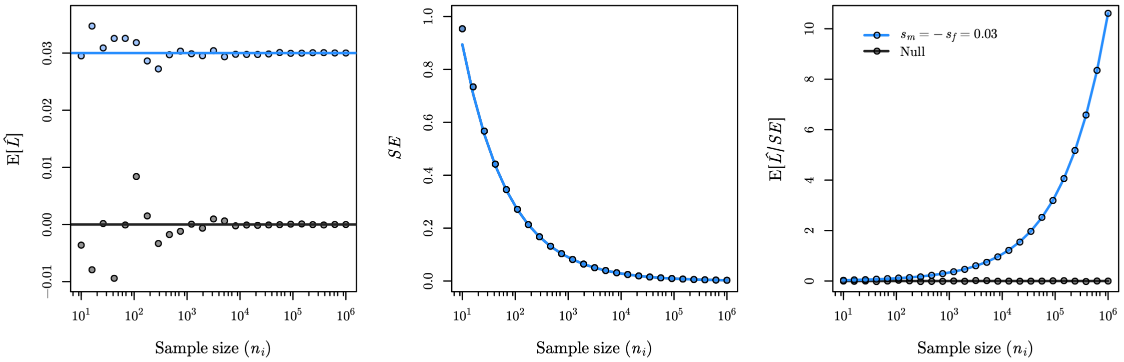


The expected distribution of the scaled sample log odds ratio will also be asymptotically normal, with

$$\frac{\hat{\mathcal{L}}}{SE} \sim \mathrm{N}\left(\frac{\log\left[\frac{(4-s_f)(4+s_m)}{(4+s_f)(4-s_m)}\right]}{8\sqrt{\frac{1}{N_f(16-s_f^2)}+\frac{1}{N_m(16-s_m^2)}}}, 1\right)$$

As noted above, the asymptotic properties of the sampling distribution of $\hat{\mathcal{L}}/SE$ break down when either MAFs are low, sample sizes are small, or both.

#### Statistical power to detect individual outlier loci

The probability that a SA locus resides within the tail of the null distribution depends on the allele frequencies at the locus, the strength of selection, and the sample size of individuals that are sequenced. The expected difference in allele frequency between the sexes, after selection, is maximized for a locus with intermediate allele frequencies and symmetric SA selection ($p$ = ½, described above), representing a best-case scenario for detecting individual outlier loci. The robustness of the normal approximation for the scaled log odds ratio greatly simplifies calculation of statistical power to identify outlier loci under sex-specific selection. For our idealized scenario where $p_f = p_m$ = ½ it is simply the area under $N\left(E\left[\frac{\hat{\mathcal{L}}}{SE}\right], 1\right)$ that falls above/below the α/2 and (1 – α)/2 quantiles of a standard normal distribution, $N(0,1)$. Below, we illustrate the probability that $\hat{\mathcal{L}}/SE$ reliably resides in the tail of the null distribution under the same parameter conditions shown for $\hat{F}_{ST}$ in Fig. 1B of the main text. Overall, the scaled log odds ratio provides similar statistical power to detect individual outlier loci as $\hat{F}_{ST}$.

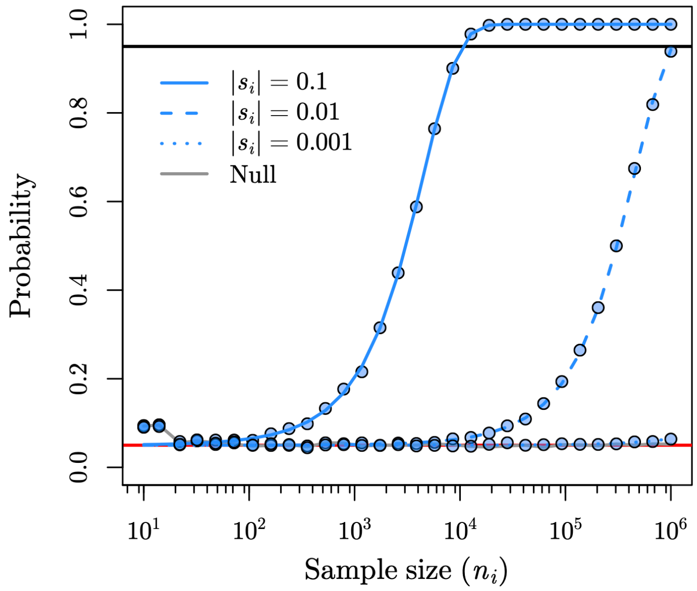


**Figure S6** Probability that $\hat{\mathcal{L}}/SE$ for an additive SA locus (*i.e.*, $h_f = h_m = ½$) with intermediate equilibrium allele frequencies ($p$, $q$ = 1/2) is in the top 2.5% tail of the null distribution $N(0{,}1)$. Sample sizes are assumed to be equal for the $i$th sex, and $|s_i|$ is the fitness cost of being homozygous for the "wrong" SA allele, where $i \in \{f{,}m\}$. Blue lines show analytic approximation based on $N\left(E\left[\frac{\hat{\mathcal{L}}}{SE}\right], 1\right)$, points show the proportion of $10^4$ simulated loci which fall in the upper and lower 2.5% tails of the null distribution. Red and black horizontal lines benchmark the 2.5% and 97.5% power thresholds.

**Multi-locus signals of sex-specific selection using $\hat{\mathcal{L}}/SE$**

Following identical logic as for $\hat{F}_{ST}$ in the main text, the full empirical distribution of $\hat{\mathcal{L}}/SE$ for autosomal SNPs may carry a cumulative signature of SA selection in the form of an inflated total number of observations in the upper quantiles of the observed $\hat{\mathcal{L}}/SE$ distribution relative to the number of observations predicted under the null. Below we illustrate the signal of enrichment of observations in the upper theoretical quantiles caused by SA selection using the ratio of observed versus permuted $\hat{\mathcal{L}}/SE$ for SNPs within 100 quantiles of the theoretical null of $\hat{\mathcal{L}}/SE$ using identical simulation methods as those presented in Fig. 2 (theory panel) of the main text. The scaled log odds ratio appears to provide a similar signal of SA selection to $\hat{F}_{ST}$, but is slightly less sensitive to shifts in the observed distribution due to SA selection (compare to Fig. 2 in the main text)

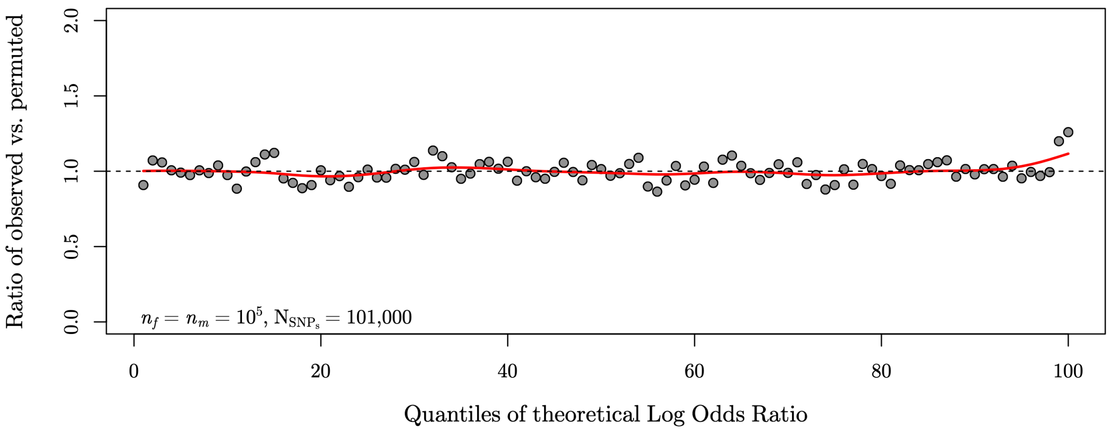


**Figure S7.** Theoretical signal of multi-locus SA polymorphism: the ratio of observed versus permuted $\hat{\mathcal{L}}/SE$ SNPs within 100 quantiles of the theoretical null of $\hat{\mathcal{L}}/SE$. **Theoretical data:** $\hat{\mathcal{L}}/SE$ values were simulated for $10^5$ neutrally evolving loci with MAF above 5% in the dataset, and for $10^3$ loci responding to SA selection prior to sampling of gene sequences (each SA-responding locus had a SA fitness effect or was in perfect linkage disequilibrium with an SA locus). SA-responding loci had intermediate allele frequencies in the population ($p$ = ½). SA selection coefficients were drawn from an exponential distribution with mean of $s_{avg}$ = 0.03, and additive fitness effects ($h$ = ½). The top 1% theoretical quantile is enriched by ~40%, implying that ~3/10 of SNPs above the 99% threshold of the null are true positives.
